# An exotic allele of barley *EARLY FLOWERING 3* contributes to developmental plasticity at elevated temperatures

**DOI:** 10.1101/2022.08.15.503967

**Authors:** Zihao Zhu, Finn Esche, Steve Babben, Jana Trenner, Albrecht Serfling, Klaus Pillen, Andreas Maurer, Marcel Quint

## Abstract

Increase in ambient temperatures caused by climate change affects various morphological and developmental traits of plants, threatening crop yield stability. In the model plant *Arabidopsis thaliana, EARLY FLOWERING 3* (*ELF3*) plays prominent roles in temperature sensing and thermomorphogenesis signal transduction. However, how crop species respond to elevated temperatures is poorly understood. Here, we show that the barley ortholog of *AtELF3* interacts with high temperature to control growth and development. We used heterogeneous inbred family (HIF) pairs generated from a segregating mapping population and systematically studied the role of exotic *ELF3* variants in barley temperature responses. An exotic *ELF3* allele from Syrian origin promoted elongation growth in barley at elevated temperatures, whereas plant area and estimated biomass were drastically reduced, resulting in an open canopy architecture. The same allele accelerated inflorescence development at high temperature, which correlated with early transcriptional induction of MADS-box floral identity genes *BM3* and *BM8*. Consequently, barley plants carrying the exotic *ELF3* allele displayed stable total grain number and mitigated yield loss at elevated temperatures. Our findings therefore demonstrate that exotic *ELF3* variants can contribute to phenotypic and developmental acclimation to elevated temperatures, providing a stimulus for breeding of climate resilient crops.

**Highlight:** We demonstrate that an exotic allele of *EARLY FLOWERING 3* (*ELF3*) contributes to plant architectural and developmental acclimation, and thereby improves yield stability at high ambient temperatures.

## Introduction

Climate change has unequivocally disrupted human and natural systems. With the global temperature predicted to reach a 1.5°C increase in the near term, global warming poses significant challenges to ecosystems and food security. The agricultural responses to global warming are of great concerns, as elevated ambient temperatures limit the productivity and spatial distribution of crop plants, resulting in severe yield losses (Leng and Huang, 2017; Zhao et al., 2017). Understanding how plants respond to elevated temperatures is pivotal to achieve crop-level adaptations and yield stability in the context of global climate change (Challinor et al., 2014; Chen et al., 2022).

Plants are constantly exposed to changing environments and are capable to adapt their growth and development accordingly. Morphological and developmental adjustments of plants induced by elevated ambient temperatures are collectively termed thermomorphogenesis (Quint et al., 2016). Our understanding of molecular genetic processes underlying thermomorphogenesis is mainly based on studies in the model plant *Arabidopsis thaliana*. In Arabidopsis, changes in ambient temperature are perceived via multiple systems. The photoreceptor phytochrome B (phyB) was the first identified plant temperature sensor, integrating light and temperature sensing (Jung et al., 2016; Legris et al., 2016). Warm temperatures accelerate the thermal/dark reversion of phyB from its active to its inactive form, thereby relieving the repression of the basic helix-loop-helix (bHLH) transcription factor PHYTOCHROME INTERACTING FACTOR 4 (PIF4) (reviewed in Delker et al., 2017). As a central regulatory hub of thermomorphogenesis signalling, PIF4 activates auxin biosynthesis genes, mediating elongation growth and leaf hyponasty at warm temperatures to improve the transpirational cooling capacity (Koini et al., 2009; Franklin et al., 2011; Crawford et al., 2012; Park et al., 2019). On the other hand, PIF4 mediates thermal acceleration of flowering by activating *FLOWERING LOCUS T (FT*) (Kumar et al., 2012). In addition to the phyB-PIF4-auxin signalling module, recent studies revealed prominent roles of the circadian clock component EARLY FLOWERING 3 (ELF3) in the Arabidopsis thermomorphogenic pathway (Box et al., 2015; Raschke et al., 2015). As an integral part of the plant circadian oscillator, the scaffold protein ELF3 interacts with LUX ARRYTHMO (LUX) and ELF4 to form the evening complex (EC) (Nusinow et al., 2011; Herrero et al., 2012). The EC directly represses the transcription of circadian clock components, as well as clock output genes such as *PIF4* (Huang and Nusinow, 2016). The prion-like domain of ELF3 functions as a thermosensor and enables the phase separation of ELF3 into liquid droplets at high temperatures, relieving the transcriptional repression of *PIF4* and potentially the direct interaction with PIF4 (Ezer et al., 2017; Jung et al., 2020). Consequently, ELF3 integrates temperature signals to the circadian oscillator, controlling rhythmic thermoresponsive outputs (Zhu et al., 2022). However, although *ELF3* homologs have been identified in a wide range of crop species (Faure et al., 2012; Zakhrabekova et al., 2012; Ning et al., 2015; Alvarez et al., 2016; Lu et al., 2017; Ridge et al., 2017), signalling components and regulation of thermomorphogenesis are largely unknown in non-model species.

As a globally cultivated crop, barley (*Hordeum vulgare* L.) has emerged as an excellent model for understanding crop adaptation to elevated temperatures (Dawson et al., 2015). The current understanding of barley temperature responses is mainly restricted to the temperature effects on reproductive development. The barley ortholog of *AtELF3* (also known as *EARLY MATURITY 8* or *Praematurum-a*) controls flowering time, and plays a critical role in barley adaptation to short growing seasons (Faure et al., 2012; Zakhrabekova et al., 2012). In barley, loss of *ELF3* function facilitates the expression of the photoperiod response gene *PHOTOPERIOD H1 (Ppd-H1*), which corresponds to Arabidopsis circadian clock gene *PSEUDO RESPONSE REGULATOR 37 (PRR37*) (Turner et al., 2005; Faure et al., 2012). Up-regulation of *Ppd-H1* promotes the expression of *FT1*, a barley homolog of Arabidopsis *FT* (Campoli et al., 2012). Induction of *FT1* is associated with the expression of barley floral meristem identity genes *VERNALIZATION 1 (VRN1), BARLEY MADS-box 3 (BM3*), and *BM8*, promoting inflorescence development (Trevaskis et al., 2007; Li et al., 2015). Alternatively, the loss of *ELF3* function mediates gibberellin (GA) biosynthesis, which cooperates with *FT1* in the acceleration of flowering (Boden et al., 2014).

Under high ambient temperatures, the floral development of barley is accelerated in an *ELF3* loss-of-function background, even with reduced or invariable *FT1* transcript levels, suggesting the involvement of *FT1*-independent pathways (Hemming et al., 2012; Ejaz and von Korff, 2017). For example, as a core component in the barley circadian network, ELF3 controls the thermo-responsiveness of *GIGANTEA* (*GI*) and *PRR* genes (Ford et al., 2016; Müller et al., 2020). Moreover, acting downstream of *ELF3*, the effects of *Ppd-H1* in thermoresponsive flowering depend on the allelic variation for *VRN1*, indicating the interaction with vernalization pathways (Ejaz and von Korff, 2017; Ochagavía et al., 2021). These findings suggest critical upstream roles of ELF3 function in the genetic control of barley developmental responses to elevated temperatures.

In addition to induced loss of function mutants, exploiting natural variation provides another approach to improve crop adaptation. Barley germplasm consists of genetically diverse landraces and exotic accessions, a large proportion of which is presumed to be adapted to various abiotic stresses (Newton et al., 2011; Russell et al., 2011). In the context of *ELF3*, for instance, a spontaneous recessive *elf3* mutant was identified in barley landraces from the Qinghai-Tibetan Plateau, where the growing season is restricted by low temperatures (Xia et al., 2017). To investigate the potential value of exotic barley accessions, the nested association mapping (NAM) population Halle Exotic Barley-25 (HEB-25) was generated by crossing 25 wild barley accessions with the elite cultivar Barke (Maurer et al., 2015). Aiming to study the effects of exotic *ELF3* variants, heterogeneous inbred family (HIF) pairs were generated based on HEB-25 (Zahn et al., 2022, Preprint). In each HIF pair, two nearly isogenic sister lines differ only in the homozygous cultivated (elite) or exotic (wild) *ELF3* allele, allowing direct comparison. Our recent findings from field experiments revealed significant roles of exotic *ELF3* alleles in barley development and grain yield, validating the previous quantitative trait locus (QTL) analysis (Maurer et al., 2016; Zahn et al., 2022, Preprint).

In this study, we systematically investigate the role of exotic *ELF3* alleles in barley temperature responses during seedling establishment, vegetative growth, and reproductive development. We demonstrate that at least one exotic *ELF3* allele interacts with elevated temperatures to modulate growth, development, and grain yield of barley.

## Materials and methods

### Plant Materials and growth conditions

We selected three HIF pairs (10_190, 16_105 and 17_041) generated from the two-rowed spring barley NAM population HEB-25, with exotic barley accessions collected from Syria, Afghanistan, and Iran, respectively (Maurer et al., 2015; Zahn et al., 2022, Preprint). The genomic setup of these three HIF pairs were previously described and is visualized in Supplementary Fig. S1 (Anand and Rodriguez Lopez, 2022; Zahn et al., 2022, Preprint). Bowman and *elf3^BW290^* (*elf3/eam8.w* loss-of-function mutant BW290 in Bowman background) were used as controls.

For the growth and development experiment, seeds were directly sown in soil and placed in growth chambers with day/night temperatures of 20°C/16°C, light intensity of 300 μmol m^-2^s^-1^ and long day photoperiod (LD; 16 h light/8 h dark). Five days after sowing (DAS), uniformly germinated seedlings were selected, and then either shifted to high ambient day/night temperatures (28°C/24°C, hereafter called 28°C treatment) or were kept at the 20°C/16°C temperature regime (hereafter called 20°C treatment). The position of plants in the phytochambers was randomly rotated twice per week. Eleven plants equaling eleven biological replicates from each genotype were used in each temperature condition.

The yield experiment was conducted at Julius Kühn-Institut (Quedlinburg, Germany). Seeds of the mutant *elf3^BW290^*, its corresponding cultivar Bowman, and HIF pair 10_190 (hereafter called HIF pair 10) seeds were coated with Rubin TT (BASF, Ludwigshafen, Germany) to avoid fungal infections before being sown in soil. Plants were grown in a greenhouse under the 20°C treatment, light intensity of 300 μmol m^-2^s^-1^ and LD (16 h light/8 h dark). Plants that reached BBCH-13 (Lancashire et al., 1991) were either shifted to the 28°C treatment conditions or were kept under the 20°C treatment. Twelve to fifteen plants from each genotype were used in each temperature condition.

### Temperature assay on plates

Bowman and *elf3^BW290^* seeds were sterilized with 4% NaClO for 30 min and were cold stratified at 4°C for 2 d in darkness. Seeds were sown on solid *Arabidopsis thaliana* solution (ATS) nutrient medium with 1% (w/v) sucrose (Lincoln et al., 1990). Vertically oriented plates were placed in darkness at 20°C to allow germination. After germination, seedlings were shifted to constant 28°C or were kept at constant 20°C, with LD (16 h light/8 h dark) and light intensity of 90 μmol m^-2^s^-1^. Leaf tips were marked after germination and seedlings were imaged in two consecutive days at Zeitgeber time (ZT) 08. The leaf length (from the marked position) was measured using RootDetection 0.1.2 (http://www.labutils.de/rd.html). Nine to seventeen seedlings from each genotype were used in each temperature condition.

### Construction of image-based phenotyping platform

To non-destructively acquire plant growth phenotypes, we constructed an image-based phenotyping platform. We customized a 1.2 m × 1.2 m × 1.8 m phenotyping frame with a stand positioned in the middle of the frame to fix the position and direction of the plants during each imaging time point. The platform included three Raspberry Pi 3 model B single-board microcomputers and three Raspberry Pi RGB camera modules (V2 8MP, Raspberry Pi foundation, Cambridge, England, UK), which enabled imaging from three directions: two side-views separated by 90° and a top-view. The lighting was provided by two light-emitting diodes (LED) lamps (4.6W, Philips, Eindhoven, Netherlands) from the top, and two LED light sets (10W, Neewer, Shenzhen, China) from the sides. The camera modules and light sets were positioned on the top of the frame or outside of it using tripods. The lens-pot distances were 1.25 m to the middle of the pot for side-views and 1.3 m to the top of the pot surface for top-view. To distinguish the plant parts from the surroundings, white foils (Colormatt-Hintergrund Super White, Studioexpress Vertriebs GmbH, Wiernsheim, Germany) were used as imaging background, blue cages (HNP Metalltechnik GmbH, Quedlinburg, Germany) were added to all pots, and blue meshes (Klartext Wunderlich Coating GmbH & Co. KG, Osterode am Harz, Germany) were used to cover the soil surface from 8 DAS.

The Raspberry Pi operation protocol for simultaneously acquiring plant images from several angles was previously described (Tovar et al., 2018). To use Windows computer as a remote host for Raspberry Pi, we used secure copy protocol (scp) to copy the public secure shell (SSH) key, and the rsync package was installed using Cygwin (https://www.cygwin.com) for proper synchronization of images.

### Phenotyping

For the growth and development experiment, plant height was manually measured without straightening the plants on a daily basis between 5 DAS and 14 DAS. From 8 DAS until 52 DAS, besides the manual measurement of plant height, total tiller number was counted, and each plant was imaged every two to four days. The images were analyzed with the HTPheno pipeline (Hartmann et al., 2011) for plant height (two side-views) and plant area (all three views). The plant volume was calculated as the square root of the product of two side-view areas and the top-view area. For convex hull area, the plant silhouette images derived from the HTPheno pipeline were further analyzed using the Hull and Circle plugin (Karperien, A., version 2.0a) in ImageJ (http://image.nih.gov/ij/). The days until visible third leaf, coleoptile tiller, first tiller, flag leaf sheath opening, and heading (first visible awns) were scored on a daily basis. The chlorophyll content was measured using the SPAD-502Plus chlorophyll meter (Konica Minolta, Chiyoda City, Tokyo, Japan) every week from 16 DAS until 52 DAS. The measurement took place at the same position: around 3 cm from the leaf collar of the second leaf. For destructive measurement of leaf size, plants (*n*=5) were harvested and the first and second leaves were imaged 16 DAS. The leaf length and width of the first and second leaves were measured using ImageJ (http://image.nih.gov/ij/). The measurement of leaf width took place at 2 cm from the leaf collar. All measurements and imaging were conducted between ZT03 and ZT06 on each measurement day.

For the yield experiment, the developmental stage was scored at 73 DAS. Plant height and total tiller number were scored at 118 DAS. At maturity (166 DAS for 28°C and 195 DAS for 20°C), the arial part of the plants was harvested. From three randomly selected spikes of each plant, the number of grains and florets per spike was determined, and the length of the spike (excluding the awns) was measured. The averaged data of each plant were used for further analysis. Plant dry weight was measured after placing plant material (arial parts excluding the spikes) into a drying oven for 1 d at 60°C. The grains were threshed using the LD180 laboratory thresher (Wintersteiger, Ried im Innkreis, Austria). The number of grains per plant, grain area, thousand grain weight, and grain yield per plant were measured using MARViN ProLine seed analyzer (MARViTECH GmbH, Wittenburg, Germany).

### Analysis of transcript levels

Plants from HIF pair 10_190 (10_elite and 10_wild) were grown under conditions as described above for the growth and development experiment. The temperature treatments started at 5 DAS. Leaf samples were harvested at ZT08 at 5, 12, 19, 27, 33, and 40 DAS, with three biological replicates each.

Total RNA was extracted using the NucleoSpin RNA Plant Kit (Macherey-Nagel), cDNA was synthesized using the PrimeScript RT Reagent Kit (Perfect Real Time, Takara Bio), and quantitative real-time-PCR was performed on an AriaMx Real-Time PCR System (Agilent) using Absolute Blue Low Rox Mix (Thermo Fisher Scientific). The expression of target genes was normalized to *HvACTIN* and *HvGAPDH* (Kikuchi et al., 2012; Zakhrabekova et al., 2012). Primer sequences are described in Supplementary Table S1.

### Analysis of *ELF3* sequence in HIF pairs

The *ELF3* coding sequences of HIF parental lines were described previously (Zahn et al., 2022, Preprint). In this study, the entire *ELF3* genomic sequence present in three HIF pairs was amplified using Ex Taq DNA Polymerase (Takara Bio, Kusatu, Shiga, Japan) and 1186 bp of promoter sequence upstream of the *ELF3* start codon in HIF pair 10 was amplified using ALLin™ RPH Polymerase (highQu GmbH, Kraichtal, Germany). The amplicons were submitted to Eurofins Genomics (Ebersberg, Germany) for dideoxy sequencing. The PCR and sequencing primers used for genomic sequence have been described previously (Zahn et al., 2022, Preprint) and those used for promoter sequence are shown in Supplementary Table S1. The sequences were aligned and compared to their corresponding parental lines (Zahn et al., 2022, Preprint) as well as their corresponding sister line in each HIF pair using MAFFT version 7 (Katoh et al., 2019).

### Growth curve modeling

Growth curves were modeled for all traits which were obtained on successive days during growth and development experiment, as cubic splines in SAS PROC TRANSREG (SAS Institute, Inc., Cary, USA) with nknots set 1. Based on the predicted values the following parameters were derived for each line: maximum increase (difference between two consecutive days), day of maximum increase, maximum value, day of maximum value, end point value, and total area under the curve (based on the trapezoidal rule with trapezoids defined between each two consecutive days).

### Principal component analysis (PCA)

PCA was performed with SAS PROC PRINCOMP (SAS Institute, Inc., Cary, USA) based on the arithmetic means of all obtained traits from the growth and development experiment, or the derived growth curve modeling traits, for each line. Due to the different units and scales of the traits, the PCA was based on the correlation matrix.

### Pairwise correlation analysis of traits

Pairwise correlation coefficients were determined for lines Bowman, *elf3^BW290^*, and HIF pair 10 between all investigated traits obtained from both growth and development, and yield experiments. As for PCA, correlations across all lines were based on the arithmetic means, as well as the derived growth curve modeling traits. All Pearson correlation coefficients and their *P* values were computed using SAS PROC CORR (SAS Institute, Inc., Cary, USA).

### Statistical analysis

Significant differences between temperature treatments and genotypes were analyzed by two-way ANOVA followed by Tukey’s HSD post-hoc test in R (R Core Team, 2013). Significant differences between two time points were analyzed by a two-sided Student’s *t*-test using GraphPad Quick-Calcs (http://graphpad.com/quickcalcs/). Linear regression was calculated and plotted using the stat_poly_eq function from the R package ggpmisc (Aphalo, 2016).

## Results

### *ELF3* is involved in barley thermomorphogenesis

In the model plant *Arabidopsis thaliana*, elevated temperatures induce a series of morphological and architectural changes, with hypocotyl elongation as a classic phenotype to represent thermo-responsiveness (Quint et al., 2016). The Arabidopsis *elf3* mutants generally display exaggerated hypocotyl growth even at relatively low temperatures, possibly due to lack of thermosensing functions (Box et al., 2015; Raschke et al., 2015; Jung et al., 2020; Zhu et al., 2022). To assess the possible role of *ELF3* in barley thermomorphogenesis, we first performed a temperature assay on agar plates mimicking the standard growth conditions for Arabidopsis seedling assays. Leaf length was measured of cultivar Bowman and *elf3^BW290^* mutant seedlings grown in LD at constant 20°C or 28°C. In two consecutive days after germination, we observed no difference in leaf length between Bowman and *elf3^BW290^* at 20°C (Fig. 1). However, at 28°C *elf3^BW290^* displayed longer leaves, compared to Bowman (Fig. 1). This suggests that *ELF3* is involved in the early temperature response of barley seedlings, controlling elongation growth in a negative manner similar to the role of Arabidopsis *ELF3* (Box et al., 2015; Raschke et al., 2015).

**Figure 1.**
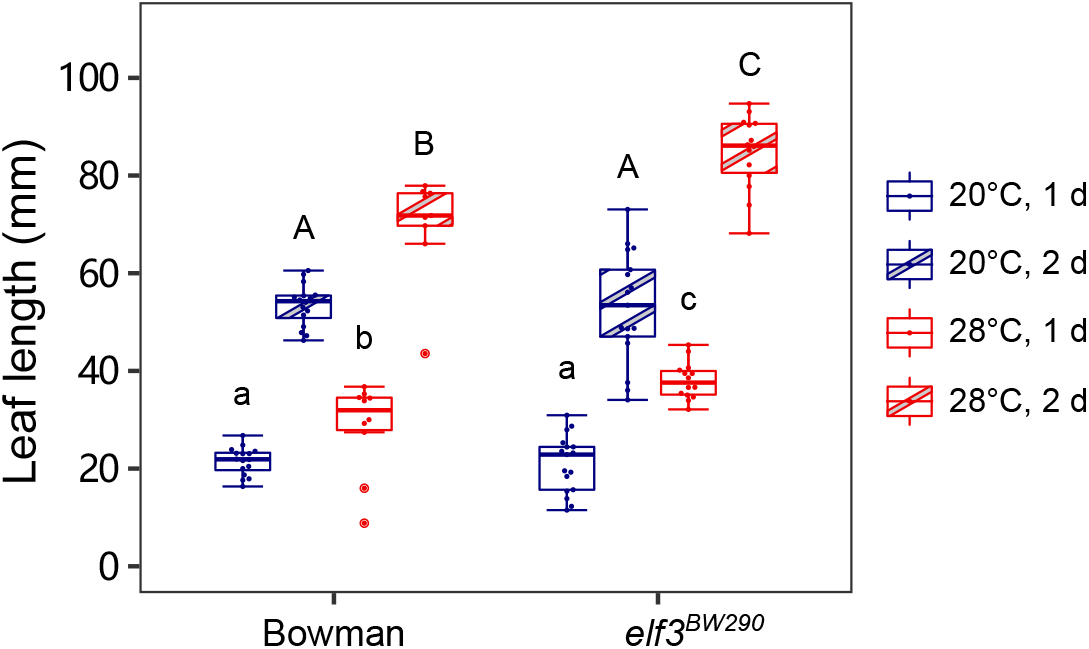
*ELF3* is involved in the temperature response of barley seedlings. Bowman and *elf3^BW290^* seedlings were grown at constant 20 or 28°C in LD (16 h light/ 8 h dark) with light intensity of 90 μmol m^-2^s^-1^. Leaf tips were marked after germination and the leaf length from the marked position was measured in two consecutive days. Boxes show medians and interquartile ranges. The whiskers extend to 1.5x interquartile ranges. Dots represent values of individual plants (*n*=9-17). Different letters above the boxes indicate significant differences (small and capital letters for the first and second days, respectively, two-way ANOVA and Tukey’s HSD test, *P*<0.05).

To further study the role of *ELF3* in barley temperature response, in addition to Bowman and *elf3^BW290^*, we selected three HIF pairs (10_190, 16_105 and 17_041) from the barley NAM population HEB-25 (Maurer et al., 2015; Maurer et al., 2016), which displayed differences for several developmental phenotypes between the HIF sister lines in previous field experiments (Zahn et al., 2022, Preprint). Each HIF pair contains two near-isogenic lines (NILs), carrying either *HvELF3* (elite) from cultivar Barke or *HspELF3* (wild, *Hsp* = *Hordeum vulgare* L. ssp. *spontaneum*) from exotic barley accessions (Supplementary Fig. S1). In this study, we initially determined the full-length genomic sequence of *ELF3* in the three used HIF pairs and compared them to the previously sequenced parental lines (Zahn et al., 2022, Preprint). We observed a W669G substitution in all three wild lines. Furthermore, the *16_105_HspELF3* (hereafter called 16_wild) and 17_wild HIF pairs belong to the same haplotype carrying four additional non-synonymous single-nucleotide polymorphisms (SNPs) in *ELF3* compared to 10_wild (Fig. 2). In line with the previous observation using barley landraces collected from the Qinghai-Tibetan Plateau (Xia et al., 2017), we identified SNPs and insertion-deletion mutations (Indels) especially in intron 2 of *ELF3* in 16_wild and 17_ wild (Fig. 2). However, the previously described alternative splicing mutation in intron 3 was not observed in our HIF lines (Xia et al., 2017). In our previous field experiments, we identified the strongest effects on shooting and heading within HIF pair 10, which could be validated also in controlled environments (Zahn et al., 2022, Preprint). Consistent with the phenotypes, we also observed differences in transcription levels of downstream genes *FT1* and *VRN1*, which are known to be regulated via ELF3, between 10_elite and 10_wild, without changes in *ELF3* transcript levels (Zahn et al., 2022, Preprint). Structural modeling of ELF3 protein variants predicted a potential influence of the W669G mutation on protein structure, which might be responsible for the observed phenotypic differences (Fig. 2) (Zahn et al., 2022, Preprint). Hence, although the previous field and indoor experiments were not performed in a temperature context, using these phenotypically divergent HIF pairs promised to provide a suitable genetic background to study the role of exotic *ELF3* alleles in barley temperature response. In the following, we systematically analyzed the role of *ELF3* on temperature-sensitive development in general by comparing Bowman and *elf3^BW290^*, and with a specific focus on natural variation between exotic *ELF3* alleles by comparing the sister lines of the three above mentioned HIF lines. We divided development into early seedling establishment, vegetative growth, and reproductive growth.

**Figure 2.**
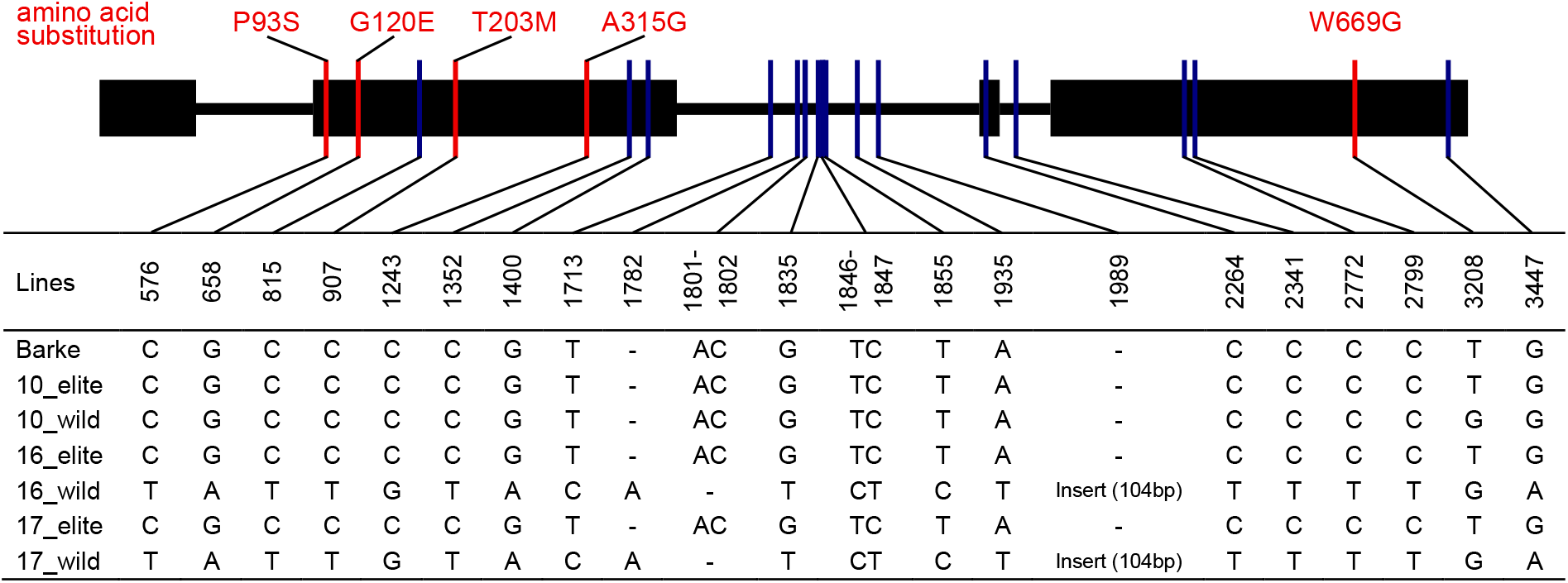
Variations in *ELF3* sequence among HIF pairs. Black rectangles (exons) and connecting lines (introns) represent the structure of barley *ELF3* (cv. Barke). Positions of the nonsynonymous single-nucleotide polymorphisms (SNPs) are shown as red vertical bars with corresponding amino acid substitutions shown above the scheme. Positions of the synonymous SNPs and the insertiondeletion mutations (Indels) in introns are shown as blue vertical bars. All mutations are listed in the table, using Barke sequence as reference. The sequence of the 104-bp-insertion at position 1989 is: AGCAAA ATGAAT GAATCT ACACTC TAAAAT ATGTCT ATATAC ATCGTA TGTAGT CCACTA GTGGAA TCTCTA GAAAGA CTTATA TTTAGG AACGGA GGGAGT AT.

### Elevated temperatures accelerate barley seedling establishment

To investigate the effect of *ELF3* and elevated temperatures on barley growth and development during early vegetative development, the eight described genotypes were grown in LD with day/ night temperatures of 20°C/16°C (20°C treatment) or 28°C/24°C (28°C treatment). The temperature treatments started 5 days after sowing (DAS), and the first phenotype, plant height, was measured on a daily basis until 14 DAS (Fig. 3A; Supplementary table S2). We observed that at elevated temperatures, plant height was significantly increased in *elf3^BW290^* from 6 DAS on. This effect was also present in the corresponding cultivar Bowman, but much delayed (reliably from 13 to 14 DAS on, Fig. 3A; Supplementary table S2). As observed in the temperature assay on plates (Fig. 1), the differences in plant height between Bowman and *elf3^BW290^* were mostly observed at 28°C but not at 20°C (Fig. 1; Fig. 3A; Supplementary table S2). Hence, plant height is obviously a phenotype that is conditionally regulated by barley *ELF3* at elevated temperature during early vegetative growth. Similar results were obtained for HIF pair 10, with 10_wild showing a stronger temperature response compared to 10_elite (Fig. 3A; Supplementary table S2), suggesting a genetic effect of the exotic barley allele on plant height during early vegetative development. In contrast, in HIF pairs 16 and 17, only temperature effects could be detected, but no allelic differences between the underlying *ELF3* variants (Fig. 3A; Supplementary table S2). Interestingly, we observed pauses of plant height increase in all genotypes during the establishment of barley seedlings, most likely because plant height was first dominated by the length of the first leaf, before the second leaf took over (Fig. 3A, B; Supplementary table S2). Notably, these pauses in plant height increase occurred earlier at elevated temperatures in almost all genotypes (Fig. 3A; Supplementary table S2), indicating accelerated growth and development.

**Figure 3.**
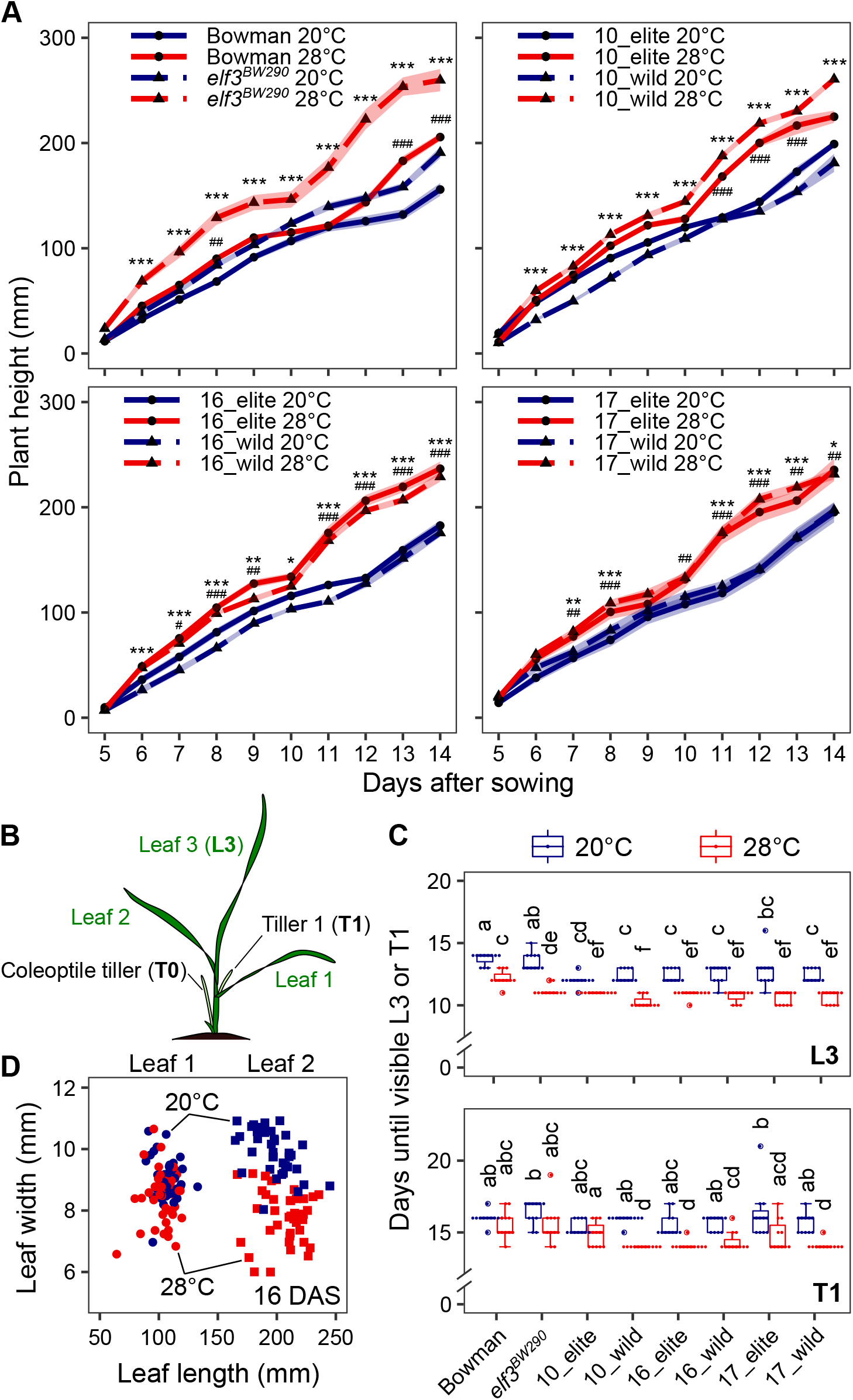
Elevated temperatures accelerate early growth and development of barley seedlings. Bowman, *elf3^BW290^* and HIF pairs were grown in LD (16 h light/ 8 h dark) at day/night temperatures of 20/16°C (20°C). At 5 days after sowing (DAS), seedlings were shifted to 28/24°C (28°C) or were kept at 20°C. (A) Plant height was measured manually on a daily basis. Lines represent the mean and ribbons indicate standard error of mean (SEM, *n*=11). Hashtags (for Bowman and HIF elite lines with *HvELF3*) and asterisks (for *elf3^BW290^* and HIF wild lines with *HspELF3*) indicate significant differences between two temperature treatments (# and *, *P*<0.05; ## and **, *P*<0.01; ### and ***, *P*<0.001; two-way ANOVA). (B) Schematic representation of a barley seedling in its three-leaf stage. (C) Days until visible third leaf (L3) and first tiller (T1). Boxes show medians and interquartile ranges. The whiskers extend to 1.5x interquartile ranges. Dots represent values of individual plants. Different letters above the boxes indicate significant differences (two-way ANOVA and Tukey’s HSD test, *P*<0.05). (D) Length and width of the first (circles) and second (squares) leaves (*n*=5) in all genotypes at 16 DAS. The genotypic effect and multiple comparisons are shown in Supplementary Table S3.

To test for accelerated development, we scored the formation of the third leaf (L3) and the first tiller (T1) under 20°C or 28°C treatment (Fig. 3B, C). In line with the results shown so far (Fig. 1; Fig. 3A; Supplementary table S2), *elf3^BW290^* displayed earlier third leaf formation compared to Bowman at 28°C but not at 20°C (Fig. 3C). In contrast, the formation of the first tiller in both Bowman and *elf3^BW290^* was neither genotype nor temperature dependent (Fig. 3C). Except for the first tiller formation in 10_elite, elevated temperatures generally accelerated the formation of the third leaf and the first tiller in HIF pairs independent of *ELF3* alleles (Fig. 3C). Prior to first tiller formation, barley seedlings can develop coleoptile tillers (T0), which arise from below ground (Fig. 3B). Coleoptile tiller development was known to be related to seedling vigor in wheat (Liang and Richards, 1994; Fujita et al., 2000), and can be suppressed by high ambient temperatures (Cannell, 1969). In confirmation of these reported observations, coleoptile tillers were largely absent at elevated temperatures, especially in Bowman and HIF pair 10 with no coleoptile tiller on 18 DAS, suggesting less vigorous seedlings despite (or maybe rather because of) accelerated growth and development (Supplementary Fig. S2).

In addition to accelerated development, elevated temperatures also cause morphological changes, for example narrower leaves in wheat (reviewed in Lippmann et al., 2019). To test whether leaf shape was influenced by temperature, we measured length and width of the first and the second leaves at 16 DAS under 20°C or 28°C treatment (Fig. 3D; Supplementary Table S3). We observed no difference in length or width in the first leaf of each genotype, possibly due to its initiation before the start of the temperature treatments (Fig. 3D; Supplementary Table S3). However, significantly narrower second leaves were observed at elevated temperatures regardless of the genotype, whereas the leaf length did not differ, suggesting reduced leaf area as previously reported (Fig. 3D; Supplementary Table S3) (Lippmann et al., 2019). Taken together, these results demonstrate that early seedling establishment is accelerated by elevated temperatures in barley, with *ELF3* alleles mainly affecting plant height.

### An exotic *ELF3* allele affects barley growth and architecture at elevated temperatures

As we observed significant effects of elevated temperatures on multiple phenotypes during barley seedling establishment (Fig. 3; Supplementary Fig. S2; Supplementary Tables S2, S3), we next asked whether the growth of barley plants would be further affected by prolonged high temperatures and whether *ELF3* alleles may differ in these responses, suggesting a functional role for *ELF3* in later vegetative growth and development. To obtain phenotypic data for barley plants in a non-destructive manner, we developed an image-based phenotyping platform based on Raspberry Pi computers and cameras (Fig. 4A) (Tovar et al., 2018). Using this platform, we were able to image plants from three directions (two side-views separated by 90° and a top-view) simultaneously (Fig. 4A). Plant traits such as plant height, plant area, and convex hull area were extracted from the obtained images (Fig. 4B). The extracted values displayed a strong correlation with the manually measured values during the experiment (Fig. 4C), demonstrating the robustness of the imaging and image analysis pipelines.

**Figure 4.**
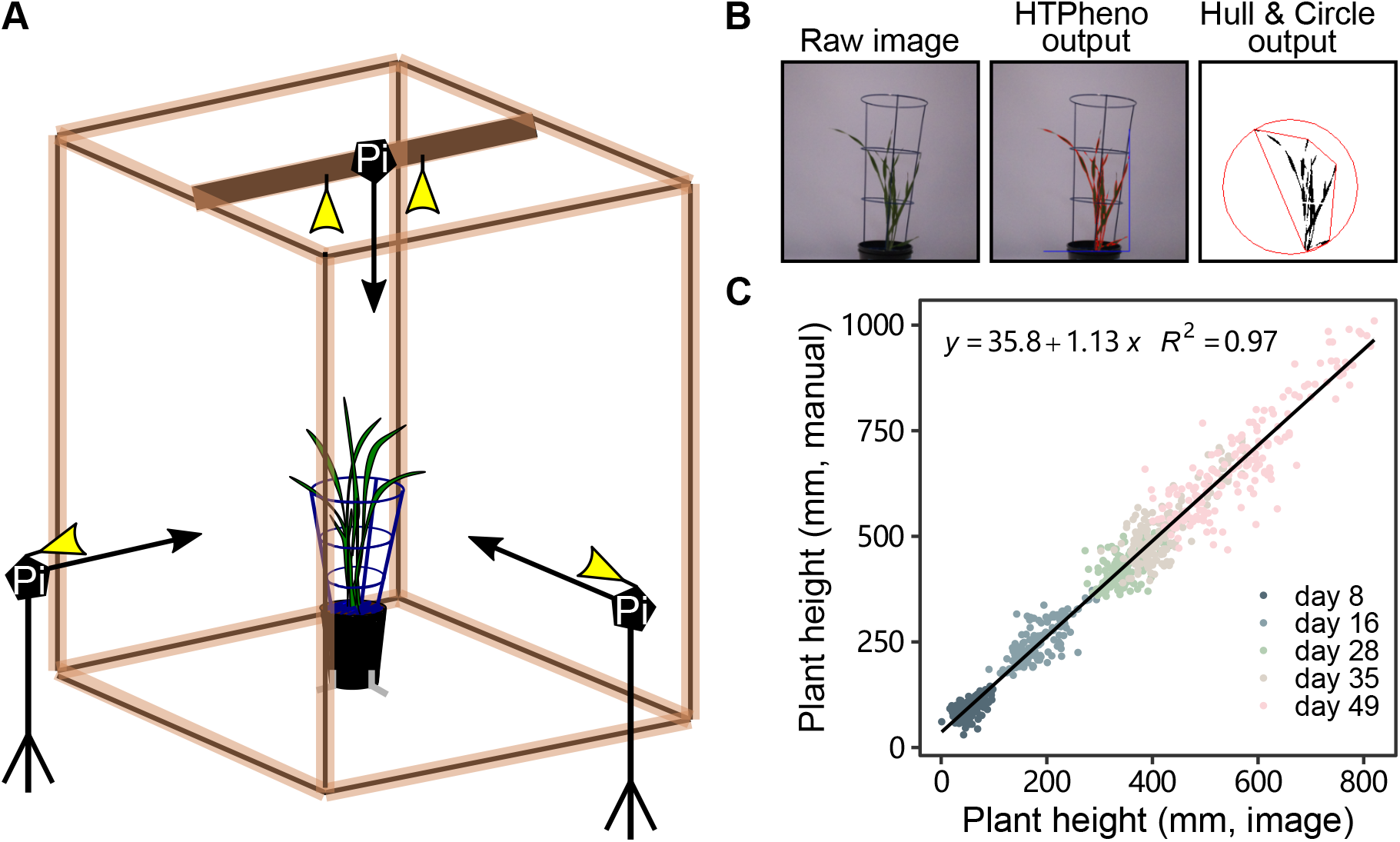
Image-based plant phenotyping for the growth and development experiment. (A) Schematic representation of the phenotyping setup. At each imaging time, images of plants were simultaneously taken from three directions. Pi, a set of a Raspberry Pi microcomputer and an RGB camera module with lighting. (B) Example of raw and output images from the image analysis pipeline. Raw images were analyzed for plant height, area, and convex hull area. Red polygon represents the convex hull with its circumcircle. (C) Linear regression of plant height values obtained from image analysis (x axis) and from manual measurement (y axis). All values (*n*=880) obtained at 8, 16, 28, 35, and 49 DAS were plotted and used for linear regression.

Starting from 8 DAS, each plant was imaged every two to four days to obtain growth curves under 20°C or 28°C treatment (Fig. 5). In confirmation of the above-described results (Fig. 3A; Supplementary Table S2), we observed a similar effect of temperature on plant height during the first two weeks of cultivation, lasting until around 30 DAS in all genotypes (Fig. 5A; Supplementary Table S4). However, after that, the positive effect of high temperature on plant height diminished and plants grown at 20°C were of similar size or even taller than those grown at 28°C (Fig. 5A; Supplementary Table S4). Although the time point when the 20°C grown plants surpassed the 28°C grown plants was genotype dependent, on 52 DAS, all genotypes displayed greater plant height at 20°C (Fig. 5A; Supplementary Table S4). These results are in line with previously reported negative effects of high temperature on plant height at maturity (Abou-Elwafa and Amein, 2016). Considering the allelic effects of *ELF3*, both the *elf3^BW290^* mutant as well as the 10_wild allele surpassed their corresponding control lines (Bowman, 10_elite) after 31 and 38 DAS, respectively, under both temperature treatments (Fig. 5A; Supplementary Table S4). In contrast, in HIF pairs 16 and 17, the elite and exotic *ELF3* alleles did not display remarkable differences in plant height, with mainly temperature effects observed (Fig. 5A; Supplementary Table S4).

**Figure 5.**
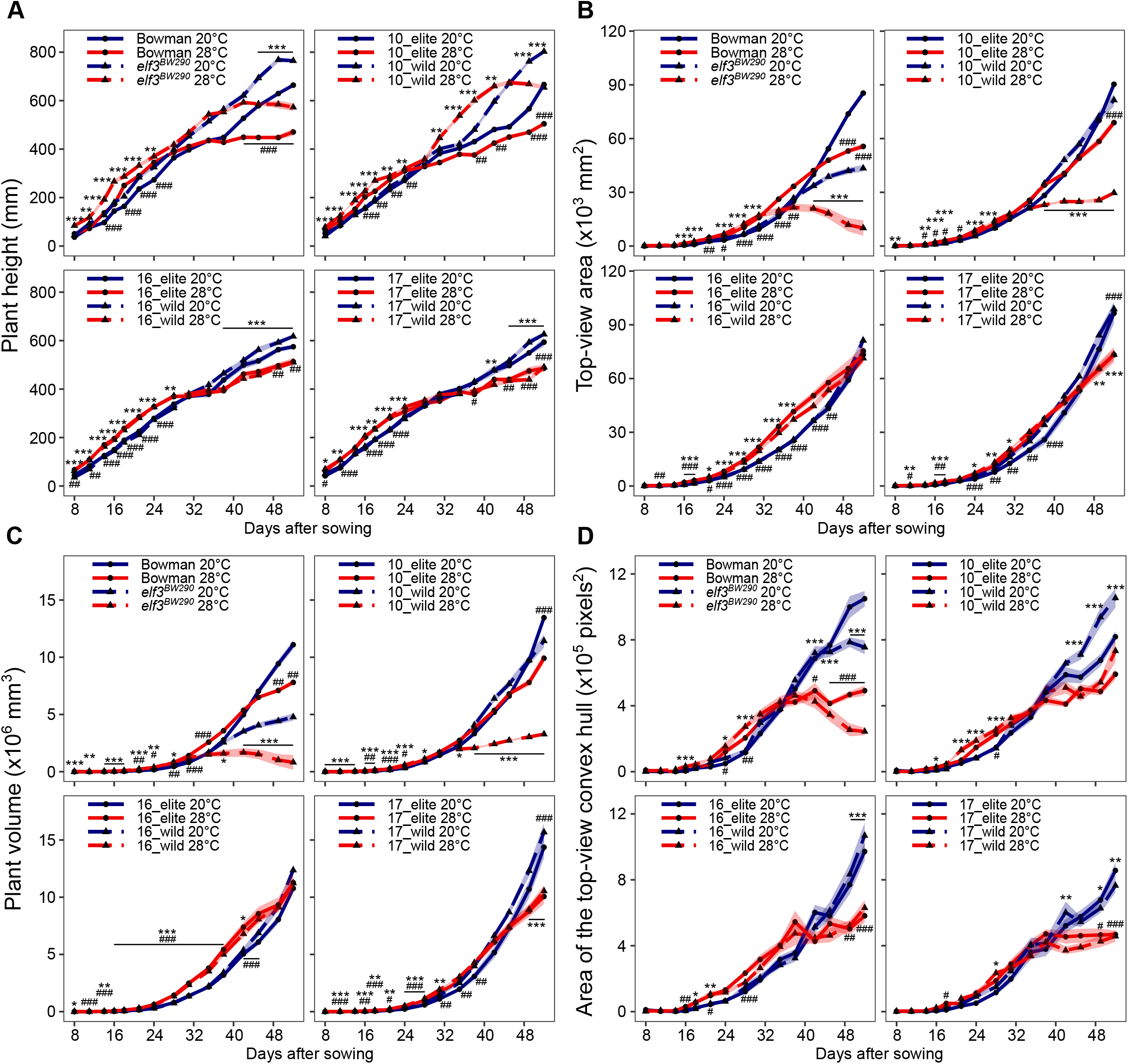
Effects of exotic *ELF3* alleles and elevated temperatures on barley growth and plant architecture. (A-D) Bowman, *elf3^BW290^* and HIF pairs were grown in LD (16 h light/ 8 h dark) at day/ night temperatures of 20/16°C (20°C). At 5 DAS, seedlings were shifted to 28/24°C (28°C) or were kept at 20°C. From 8 DAS, plants were imaged every two to four days. Plant height (A, average of two side-views) and top-view plant area (B) were obtained using HTPheno pipeline. (C) Plant volume was estimated using plant area data from top-view (B) and two side-views. (D) Top-view convex hull area was obtained using Hull & Circle pipeline. Lines represent the mean and ribbons indicate SEM (*n*=11). Hashtags (for Bowman and HIF elite lines with *HvELF3*) and asterisks (for *elf3^BW290^* and HIF wild lines with *HspELF3*) indicate significant differences between two temperature treatments (# and *, *P*<0.05; ## and **, *P*<0.01; ### and ***, *P*<0.001; two-way ANOVA).

Similar to plant height, we observed reduced plant area at elevated temperatures at 52 DAS, except for HIF pair 16 showing no difference between temperatures (Fig. 5B; Supplementary Table S5). Albeit greater in plant height, the plant area of *elf3^BW290^* was smaller than Bowman from 45 DAS at 20°C and from 35 DAS at 28°C (Fig. 5B; Supplementary Table S5). However, plant area of 10_elite and 10_wild plants did not differ at 20°C, but only at 28°C (Fig. 5B; Supplementary Table S5). In general, from 42 DAS, *elf3^BW290^* and 10_wild at 28°C displayed lowest plant areas from both top- and side-views among all genotypes (Fig. 5B; Supplementary Fig. S3A; Supplementary Tables S5, S6). As such, extended elongation growth seems to come at the cost of reduced leaf area for light interception and photosynthesis, which likely depends on *ELF3*. To represent plant biomass, the plant volume was estimated based on the plant areas of top- and two side-views, showing similar trends as plant areas (Fig. 5C; Supplementary Table S7).

Interestingly, although plant area was reduced in *elf3^BW290^* and 10_wild at 28°C, we observed relatively high convex hull areas from both top- and side views, especially in 10_wild (Fig. 5D; Supplementary Fig. S3B; Supplementary Tables S8, S9). The convex hull area (the area of the red polygon in Fig. 4B) represents the smallest area enclosing the whole plant silhouette. Different to plant area, the convex hull area is mainly contributed by leaf length and bolting, representing the expansion of plants. Using both parameters (area and convex hull area) allowed a more comprehensive description of plant architecture. Our observations of reduced plant area but increased or stable convex hull area in *elf3^BW290^* and 10_wild at 28°C could be a consequence of thinner leaves at 28°C (Fig. 3D; Supplementary Table S3). This conclusion is generally acceptable as the total tiller number on 52 DAS was not different (*elf3^BW290^*, HIF pair 10, 16_elite and 17_elite) or even higher (Bowman, 16_wild and 17_wild) at 28°C (Fig. 6A; Supplementary Table S10). The larger convex hull area is a proxy for a more openly structured habitus of the shoot. From work in Arabidopsis it is known that such loose architectural adjustments promote ventilation and thereby facilitate evaporative leaf cooling at elevated temperatures (Crawford et al., 2012).

**Figure 6.**
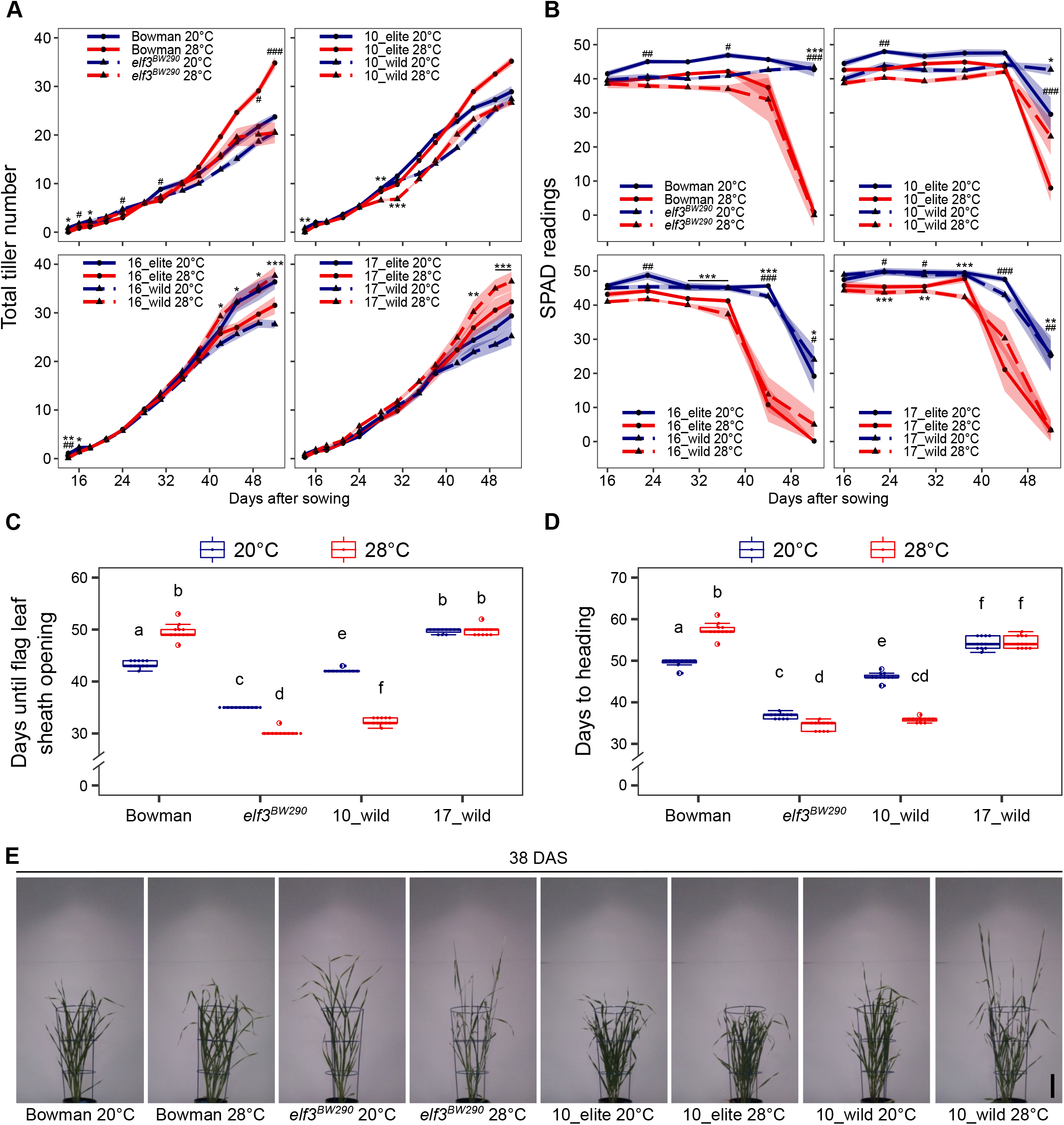
Effects of exotic *ELF3* alleles and elevated temperatures on barley tillering, leaf senescence, and heading. (A-E) Bowman, *elf3^BW290^* and HIF pairs were grown in LD (16 h light/ 8 h dark) at day/night temperatures of 20/16°C (20°C). At 5 DAS, seedlings were shifted to 28/24°C (28°C) or were kept at 20°C. (A) Total tiller number was scored from 14 DAS. (B) Chlorophyll content of the second leaf was measured every week from 16 DAS. Lines represent the mean and ribbons indicate SEM (*n*=11). Hashtags (for Bowman and HIF elite lines with *HvELF3*) and asterisks (for *elf3^BW290^* and HIF wild lines with *HspELF3*) indicate significant differences between two temperature treatments (# and *, *P*<0.05; ## and **, *P*<0.01; ### and ***, *P*<0.001; two-way ANOVA). Days until flag leaf sheath opening (C) and heading (D) were scored in the lines showing visible awns before the end of the experiment. Boxes show medians and interquartile ranges. The whiskers extend to 1.5x interquartile ranges. Dots represent values of individual plants. Different letters above the boxes indicate significant differences (two-way ANOVA and Tukey’s HSD test, *P*<0.05). (E) Representative images of Bowman, *elf3^BW290^*, 10_elite, and 10_wild plants on 38 DAS. Scale bar = 10 cm.

To assess the overall phenotypic responses to elevated temperatures during vegetative growth and development, a principal component analysis (PCA) was carried out based on the arithmetic means of all obtained traits (Supplementary Fig. S4A). The first two principal components (PC1 and PC2) accounted for 75.8% of the variance. PC1 separated samples by temperature treatment, whereas PC2 separated samples by genotype. While *elf3^BW290^* displayed a clear divergence of from other genotypes at both temperatures, this separation was observed in 10_wild only at 28°C (Supplementary Fig. S4A). To avoid a potential bias in correlation caused by large amounts of non-significant data as part of the growth curve measured during early vegetative stage, we performed an additional PCA based on the growth curve modeling traits (Supplementary Fig. S4B). Using derived modeling traits, we were able to tone down the wealth of data and focus on important time points (e.g., day of maximum increase, day of maximum value, and end point) as well as general patterns of the growth curves (e.g., total area under the curve) instead of considering all measurement time points identically in the analysis. Again, clustering of *elf3^BW290^* and 10_wild at 28°C was detected (Supplementary Fig. S4B).

Taken together, our data so far demonstrate that the Syrian *ELF3* allele in HIF pair 10 and *elf3^BW290^* mutant tend to behave similarly during vegetative growth and development. We can therefore conclude that *ELF3* in general but also naturally occurring genetic variation in wild barley populations contribute to architectural changes of shoot tissues at elevated temperatures. Similar behavior of the exotic 10_wild allele and the *ELF3* loss-of-function mutant *elf3^BW290^* (which carries a truncated ELF3 protein due to a premature STOP codon) poses the question of functionality of the encoded protein variant 10_wild, which is, however, out of the scope of this study.

### An exotic *ELF3* allele accelerates barley developmental growth at elevated temperatures

The plant life cycle can be divided into distinct phases from germination to senescence, and the timing of transition from one phase to the next can be either accelerated or delayed by high temperatures, depending on plant species and temperature regimes (reviewed in Lippmann et al., 2019). We therefore asked whether exotic *ELF3* alleles are involved in controlling developmental timing of leaf senescence and flowering at elevated temperatures. As chlorophyll degradation is one of the hallmarks of leaf senescence (Guo et al., 2021), we monitored the chlorophyll content of the second leaf every week during the extended vegetative growth and development experiment (Fig. 6B; Supplementary Table S11). As expected, we observed that all genotypes displayed earlier leaf senescence at 28°C compared to 20°C (Fig. 6B; Supplementary Table S11). Although the decrease of chlorophyll content was not *ELF3* dependent, we observed earlier onset of leaf senescence at both temperatures in HIF pairs 16 and 17, indicating an effect of the genetic background in these lines outside of the *ELF3* locus (Fig. 6B; Supplementary Table S11). Consistent with previous reports from various species (Djanaguiraman and Prasad, 2010; Lobell et al., 2012; Shirdelmoghanloo et al., 2016), our data suggest premature leaf senescence at elevated temperatures (Fig. 6B; Supplementary Table S11), but argues against a role of *ELF3* in its regulation.

Senescence of old leaves enables remobilization and retranslocation of nutrients to newly formed organs (e.g., sink leaves or seeds), and often correlates with the timing of flowering (Kim et al., 2020). To test whether and how flowering time is affected by elevated temperatures and *ELF3* alleles, we scored the days until flag leaf sheath opening (BBCH-47, Fig. 6C) and the days until heading as a proxy for flowering time (BBCH-49, Fig. 6D). In contrast to Arabidopsis, but consistent with a previous report in barley (Ejaz and von Korff, 2017), we observed that Bowman displayed delayed flag leaf sheath opening and heading at 28°C compared to 20°C, whereas *elf3^BW290^* displayed the opposite temperature effect (Fig. 6C, D). Although 10_elite plants did not finish flowering at both temperatures until the end of the experiment (62 DAS, data therefore omitted from Fig. 6C, D), 10_wild plants flowered much earlier (Fig. 6D, E). Interestingly, the early flowering of 10_wild plants was further accelerated by elevated temperatures, displaying an even larger temperature response compared to *elf3^BW290^* (Fig. 6D, E). The early flowering of *elf3^BW290^* and 10_wild at 28°C indicates early bolting, which could partially explain previously observed architectural changes of these two lines (Fig. 5; Fig. 6C-E; Supplementary Figs. S3, S4; Supplementary Tables S4-9). In addition, the flowering time of 17_wild was not temperature dependent (Fig. 6D), whereas the other genotypes (HIF pair 16 and 17_elite) did not start flowering until the end of the experiment (data therefore omitted from Fig. 6C, D). Taken together, in the vast majority of phenotypes the sister lines of HIF pair 10 differed. We therefore focused further analyses on these genotypes.

To better understand the differences in the timing of vegetative to reproductive phase transition between the elite and wild alleles of HIF pair 10, we first assessed the allelic effects on barley meristem development and found that from 19 DAS, 10_wild plants displayed faster inflorescence devel-opment than 10_elite at both temperatures (Fig. 7A). In contrast to 10_elite, the meristem development of 10_wild was drastically accelerated by elevated temperatures, consistent with the flowering time results (Fig. 6C-E; Fig. 7A). To substantiate these observations on the molecular and regulatory level, we investigated the transcriptional behaviour of floral regulator genes in leaf samples from the plants used for meristem dissection (Fig. 7A, B; Supplementary Table S12). With few exceptions, transcript abundance of *Ppd-H1, FT1*, and *VRN1* remained largely unaltered by temperature or *ELF3* allele (Fig. 7B; Supplementary Table S12). The exceptions were that 10_elite at 28°C had reduced transcript levels of *Ppd-H1* (19 to 27 DAS) and *VRN1* (27 DAS) compared to 20°C, and 10_wild at 28°C displayed reduced transcript levels of *Ppd-H1* and *FT1* on 40 DAS (Fig. 7B; Supplementary Table S12). During meristem development, transcript abundance of MADS-box genes *BM3* and *BM8* increased in both lines and temperature conditions (Fig. 7B; Supplementary Table S12). However, the onset of *BM3* and *BM8* expression occurred already at 19 DAS in 10_wild at 28°C, which is earlier compared to 10_wild at 20°C and 10_elite (Fig. 7B; Supplementary Table S12). Correlated with flowering time and meristem development (Fig. 6C-E; Fig. 7A), 10_wild plants displayed induced expression of *BM3* and *BM8* at 28°C between 19 to 33 DAS, when compared to 10_elite under the same conditions (for *BM3*), or 10_wild at 20°C (for *BM8*) (Fig. 7B; Supplementary Table S12), in line with the time points displaying morphological differences in shoot apical meristems displayed in Fig. 7A. Importantly, across development *ELF3* transcript levels hardly varied between both alleles in the vast majority of time points (Fig. 7B; Supplementary Table S12). Moreover, ~1.2 kb of promoter sequence upstream of the *ELF3* start codon was identical between both alleles (Supplementary Dataset S1), suggesting that the observed phenotypes including the differences in transcript abundance of downstream target genes are due to post-transcriptional differences between both alleles, possibly on the level of functional protein.

**Figure 7.**
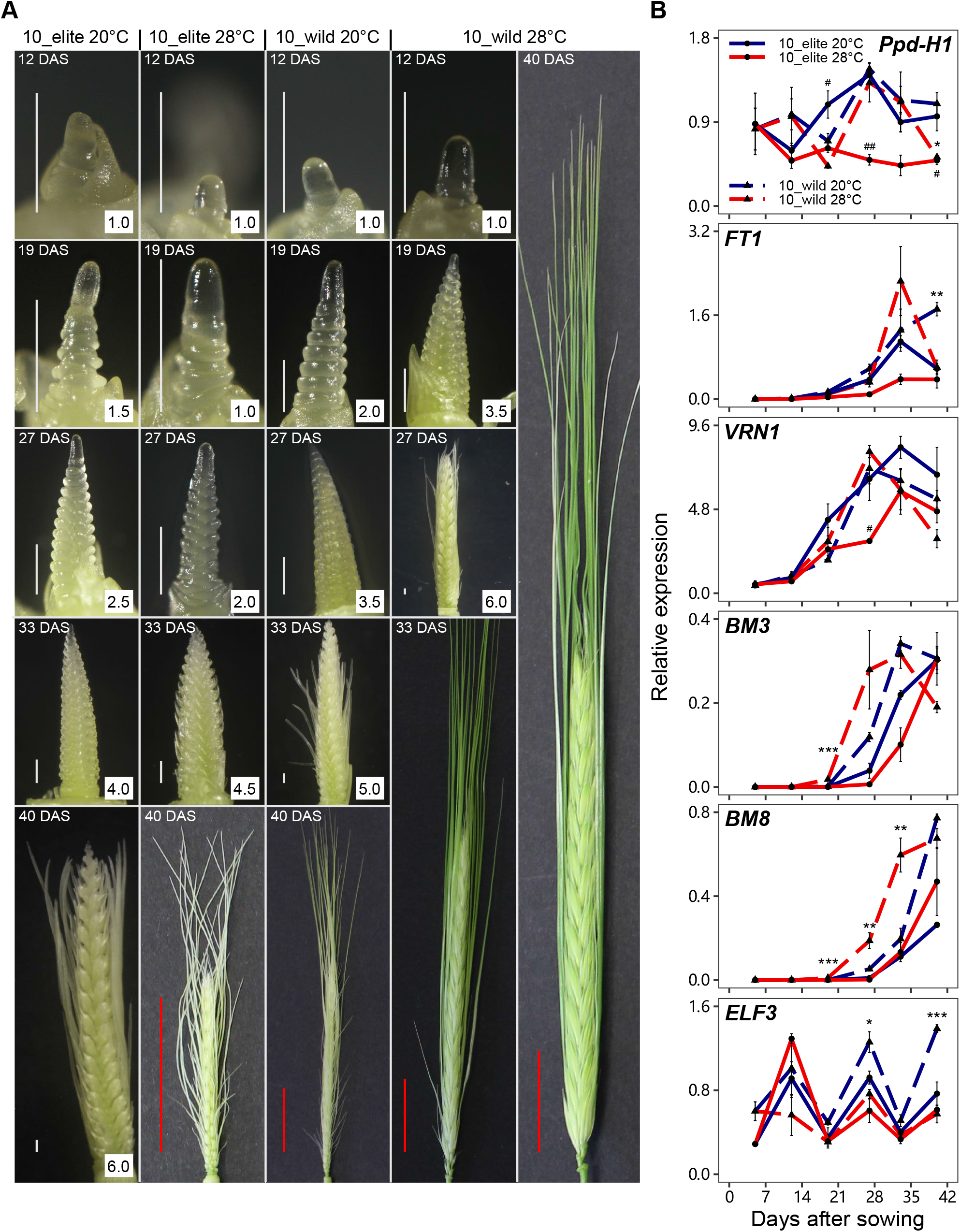
An exotic *ELF3* allele interacts with elevated temperatures to control meristem development and the transcript levels of flowering genes. (A) Representative images of shoot apex meristem and inflorescence in 10_elite and 10_wild plants grown in LD (16 h light/ 8 h dark) at day/night temperatures of 20/16°C (20°C). At 5 DAS, seedlings were shifted to 28/24°C (28°C) or were kept at 20°C. Three plants were harvested per genotype per temperature at 12, 19, 27, 33, and 40 DAS. Figures in white boxes indicates Waddington scale scores until 6.0. Scale bars, white = 500 μm; red = 2 cm. (B) Transcript levels of barley flowering genes. Leaf samples were harvested at ZT08 during meristem dissection at each time point in (A), as well as at 5 DAS before the start of temperature treatment. Expression levels were normalized to *HvACTIN* and *HvGAPDH*. Error bars indicate the SEM (*n*=3) of three biological replicates. Hashtags (for 10_elite) and asterisks (for 10_wild) indicate significant differences between two temperature treatments (# and *, *P*<0.05; ## and **, *P*<0.01; ### and ***, *P*<0.001; two-way ANOVA).

Together, flowering time data, floral meristem dissection and gene expression analyses demonstrate that the exotic *ELF3* allele in HIF pair 10 accelerates barley developmental growth at elevated temperatures, putatively by promoting the transcript levels of floral regulators including *BM3* and *BM8* MADS-box genes.

### An exotic *ELF3* allele has reduced yield loss at elevated temperatures

Under climate change scenarios, crop production and grain yield are predicted to be severely threatened by rising temperatures (Battisti and Naylor, 2009; Asseng et al., 2015). With ambient temperature increased by 7 to 8°C, striking yield losses were observed in barley (Ejaz and von Korff, 2017; Ochagavía et al., 2021). Interestingly, the early flowering mutant *elf3^BW290^* was reported to maintain seed number at elevated temperatures (Ejaz and von Korff, 2017; Ochagavía et al., 2021), thereby representing some sort of increased yield stability. Since (i) the exotic *ELF3* allele in HIF pair 10 behaved similar to *elf3^BW290^* in many of the assays performed so far and (ii) to assess the effect of 10_wild on yield at elevated temperatures, we performed an additional experiment in the greenhouse. In this experiment, four genotypes (Bowman, *elf3^BW290^*, and HIF pair 10) were grown in LD with day/night temperatures of 20°C/16°C (20°C treatment) or 28°C/24°C (28°C treatment). To avoid the effects of elevated temperatures on early seedling establishment (Fig. 3; Supplementary Fig. S2; Supplementary Tables S2, S3), the temperature treatments started from the three-leaf stage (BBCH-13, 15 to 17 DAS) (Lancashire et al., 1991). We first examined whether the growth and developmental traits under greenhouse conditions were comparable to the results from the environmentally better controlled phytochamber experiments. Despite the expected generally delayed phase transition under greenhouse conditions, we observed early flowering in *elf3^BW290^* at both temperatures and in 10_wild at 28°C (Supplementary Fig. S5A); all four genotypes displayed reduced plant height at 28°C 118 DAS (Supplementary Fig. S5B). Total tiller number was not changed in Bowman but reduced in *elf3^BW290^* at elevated temperatures 118 DAS (Supplementary Fig. S5C). In contrast, both lines in HIF pair 10 had more tillers at 28°C than 20°C (Supplementary Fig. S5C). Taken together, although the total tiller number data were different (Fig. 6A; Supplementary Fig. S5C; Supplementary Table S10), the vast majority of phenotypes from the phytochember experiments was reproducible in the greenhouse (Figs. 5A, 6A, C-E, 7; Supplementary Fig. S5A, B; Supplementary Tables S4, S12). This suggested that documentation of yield related phenotypes in the greenhouse may as well provide reliable insight into the roles of *ELF3* and especially the exotic 10_wild allele.

At maturity, we measured plant dry weight, several spike parameters (ear length and grains/florets per spike), and several grain parameters (grain number, grain area, thousand grain weight, and grain yield) (Fig. 8; Supplementary Fig. S5D-F). We found plant dry weight to be reduced in Bowman and 10_elite at elevated temperatures, but it was not influenced by temperature in *elf3^BW290^* and 10_wild (Fig. 8A). Under both temperatures, plant dry weight was lower in *elf3^BW290^* and 10_wild when compared to Bowman and 10_elite, respectively (Fig. 8A). The ear length of Bowman and *elf3^BW290^* was not affected by temperature, with *elf3^BW290^* having shorter ears than Bowman (Supplementary Fig. S5D). Shorter ears were also observed in 10_wild compared to 10_elite, but only at elevated temperatures (Supplementary Fig. S5D). At 28°C, the number of grains and florets per spike was reduced in Bowman, 10_elite, and 10_wild, but not in *elf3^BW290^* (Fig. 8B; Supplementary Fig. S5E). The *elf3^BW290^* phenotype was consistent with the previous report from (Ejaz and von Korff, 2017). *elf3^BW290^* had less grains and florets per spike than Bowman under both temperatures, whereas 10_wild only showed reduced grain and floret numbers per spike at 28°C compared to 10_elite (Fig. 8B; Supplementary Fig. S5E). The ratio of grains and florets per spike was slightly reduced in 10_elite and 10_wild at 28°C (Supplementary Fig. S5F), indicating the negative effects of high temperature on floret fertility as previously reported for wheat (Prasad and Djanaguiraman, 2014).

**Figure 8.**
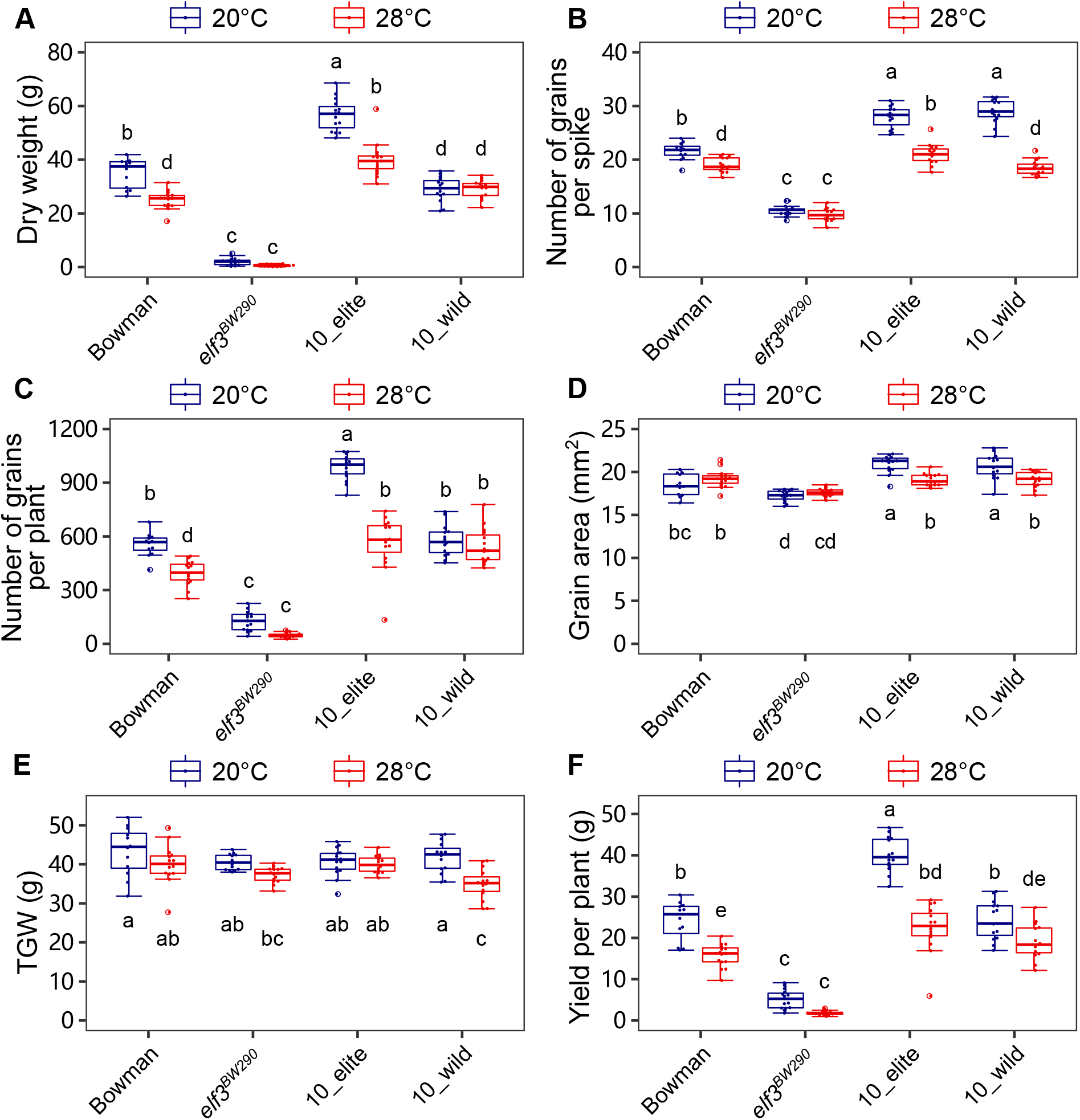
An exotic *ELF3* allele mitigates yield loss at elevated temperatures. Bowman, *elf3^BW290^*, and HIF pair 10 plants were grown in greenhouse conditions with LD (16 h light/ 8 h dark) and day/night temperatures of 20/16°C (20°C). Plants reached three-leaf stage were shifted to 28/24°C (28°C) or were kept at 20°C. At maturity, plant dry weight (A), number of grains per spike (B) and per plant (C), grain area (D), thousand grain weight (E), and yield per plant (F) were scored. Boxes show medians and interquartile ranges. The whiskers extend to 1.5x interquartile ranges. Dots represent values of individual plants (*n*=12-15). Different letters above or below the boxes indicate significant differences (two-way ANOVA and Tukey’s HSD test, *P*<0.05).

The total grain number per plant displayed similar trends as the number of grains per spike, except that 10_wild had reduced total grain number at 20°C (Fig. 8B, C). These results could be explained by reduced spike or reproductive tiller number, as the total tiller number was not different in 10_wild compared to 10_elite at 20°C (Supplementary Fig. S5C). The grain area was not affected by temperature in Bowman and *elf3^BW290^*, with *elf3^BW290^* having smaller grain area compared to Bowman (Fig. 8D). In contrast, 10_elite and 10_wild displayed the same grain area which is reduced at elevated temperatures (Fig. 8D). The thousand grain weight (TGW) was not affected by temperature in Bowman, *elf3^BW290^* and 10_elite, whereas 10_wild had reduced TGW at elevated temperatures (Fig. 8E). The reduction in TGW of 10_wild at 28°C resulted in reduced yield per plant, whereas the yield of the other lines was mostly dependent on the number of grains per plant (Fig. 8C, E, F). Albeit 10_wild displayed striking yield loss at 20°C compared to 10_elite, the yield of these two lines was not different at 28°C (Fig. 8F). These data suggest that the yield loss caused by elevated temperatures are mainly due to reduced grain number, which is mitigated in HIF pair 10 with the exotic *ELF3* allele.

To understand whether the yield parameters are linked to morphological and/or developmental traits, we analyzed putative correlations of temperature responses amongst all traits in Bowman, *elf3^BW290^*, and HIF pair 10 (Supplementary Dataset S2). Pairwise correlations of selected traits are shown in Fig. 9. As expected, high correlations can be observed among traits within similar growth and developmental stages. During early vegetative stage, temperature-induced leaf and tiller formation correlated strongly with early plant architectural traits (e.g., convex hull area, area, and volume, 16 DAS, Fig. 9). Similarly, temperature-induced reduction in yield strongly correlated with grain number, plant dry weight, as well as late plant architectural traits (e.g., area and volume, 52 DAS, Fig. 9). Interestingly, although temperature response of TGW was not correlated with any other yield parameters, it correlated positively with late plant architectural traits but negatively with early plant architectural traits (Fig. 9). These correlation patterns indicate potential connections of traits in barley response to elevated temperatures.

**Figure 9.**
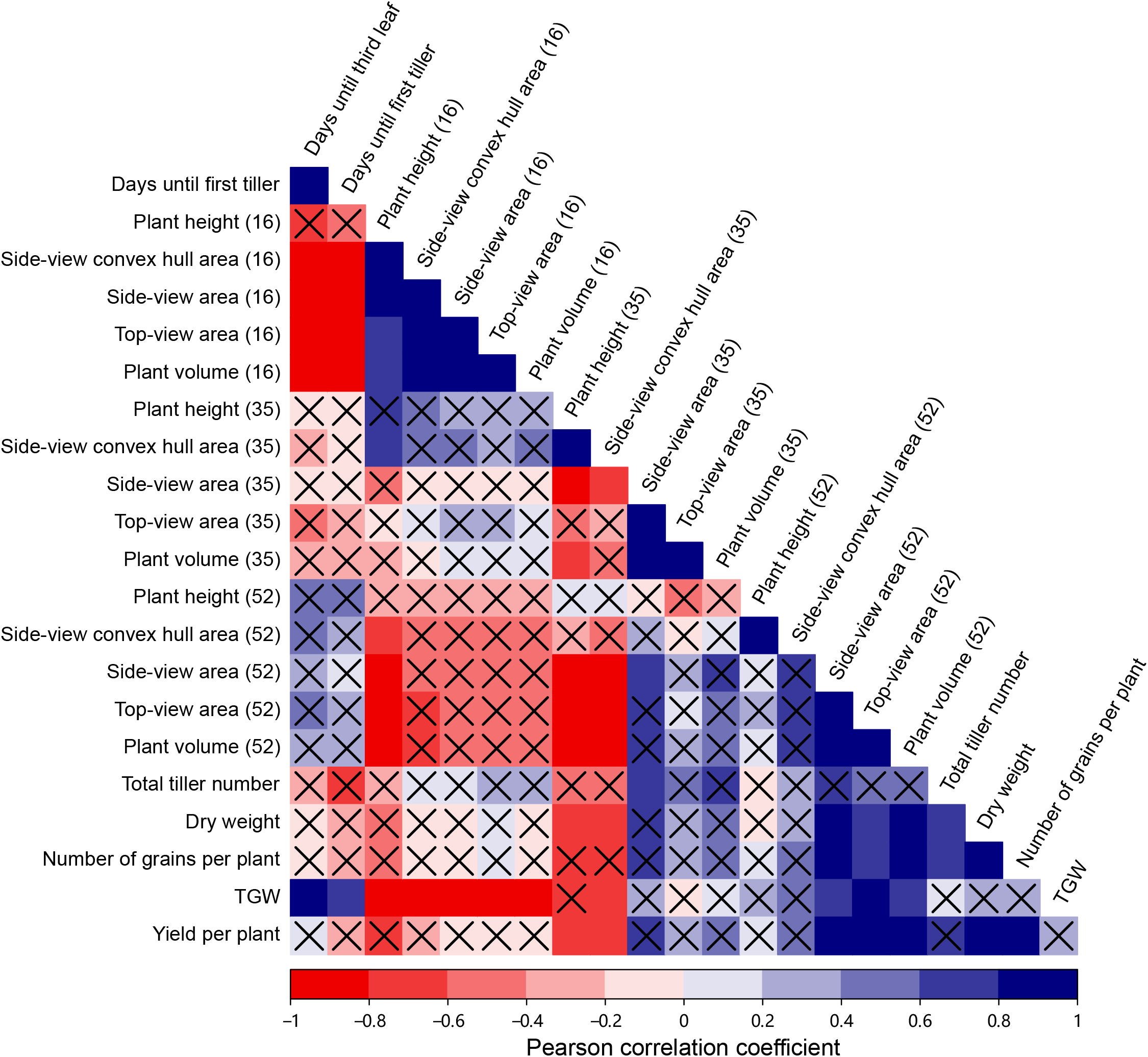
Correlation of temperature responses in selected traits. Pairwise correlation coefficients were determined for Bowman, *elf3^BW290^*, and HIF pair 10 between selected traits obtained from both growth and development, and yield experiments. Numbers in the bracket indicate DAS. Pearson correlation coefficients were tested for significance and only significant coefficients with *P*<0.05 are not crossed. Correlations for all measured and derived growth curve modeling traits are shown in Supplementary Dataset S2.

Taken together, our data demonstrate that an exotic barley *ELF3* allele not only contributes to thermomorphogenic architectural adjustments during vegetative growth, but also accelerates phase transition at elevated temperatures, mitigating further yield loss.

## Discussion

As global temperature is predicted to keep increasing in the near term, generating temperature-resilient crop varieties becomes an urgent need to avoid yield loss. Key regulators that provide phenotypic and developmental plasticity in response to elevated temperatures are poorly understood in major staple crops like barley. In this study, we systematically analyzed the effects of elevated temperatures across the barley life cycle, including seedling establishment, vegetative to reproductive stage transition, and yield at maturity. Using HIF pairs generated from a segregating mapping population with wild or elite *ELF3* alleles but an otherwise largely fixed genomic background allowed us to study the role of naturally occurring genetic variation in *ELF3* in barley thermomorphogenesis across a full life cycle. Here, we demonstrate that an exotic *ELF3* allele interacts with high ambient temperature to accelerate growth and development in barley as an acclimation response, improving yield stability.

Thermomorphogenesis as a developmental process has been neglected for a long time. But studies in the model organism Arabidopsis and elucidation of the underlying regulatory mechanisms have significantly advanced the field in the last 25 years (Gray et al., 1998; Delker et al., 2022). In the model plant, elevated temperatures induce elongation of hypocotyl, petioles, as well as leaf hyponasty (Crawford et al., 2012; Quint et al., 2016; Park et al., 2019). The barley lines used in our study likewise displayed increased plant height (Figs. 1, 3A, 5A; Supplementary Tables S2, S4) and accelerated formation of leaves and tillers (Fig. 3C), which were coupled with reduced leaf width (Fig. 3D; Supplementary Table S3) and the absence of a coleoptile tiller (Supplementary Fig. S2). As these thermoresponsive phenotypes are consistent with previous reports in barley and wheat (Cannell, 1969; Huang and Taylor, 1993; Abou-Elwafa and Amein, 2016), we conclude that the general phenotypic responses to elevated ambient temperatures during early vegetative development seem to be largely conserved across species. During seedling establishment, the *elf3^BW290^* mutant allele and the exotic allele 10_wild, which was derived from the HEB-25 mapping population (Maurer et al., 2015), mainly interacted with temperature in controlling elongation growth (Figs. 1, 3A, 5A; Supplementary Tables S2, S4), again similar to *elf3* mutant phenotypes in Arabidopsis. Via its functions as a thermosensor (through its prion-like domain) and as a core component of the circadian clock, Arabidopsis ELF3 negatively controls thermomorphogenesis via PIF4 (Box et al., 2015; Raschke et al., 2015; Jung et al., 2020; Zhu et al., 2022). As elongation of cereal leaves likely employs different mechanisms than those of dicots (Fricke, 2002) and ortholog relationships between Arabidopsis and barley *PIF*s are not phylogenetically traceable, a major regulatory hub like PIF4 in Arabidopsis is unknown in barley. Therefore, any assumption about the regulatory level of thermomorphogenesis in barley would be purely speculative.

Upon prolonged exposure to high temperature, plant height, plant area, and estimated biomass were reduced (Fig. 5A-C; Supplementary Fig. S3; Supplementary Tables S4-7). In combination with tillering and accelerated flowering, these phenotypes collectively resulted in architectural changes in the *elf3^BW290^* mutant and 10_wild (Figs. 5, 6A, C-E, 7A; Supplementary Figs. S3, S4; Supplementary Tables S4-10). Such architectural acclimation can be characterized as well-distributed and distanced plant organs (Fig. 6E), analogous to the open rosette structure in Arabidopsis, which improves leaf cooling and maintains photosynthetic efficiency at elevated temperatures (Crawford et al., 2012; Park et al., 2019). Interestingly, *elf3^BW290^* displayed such architecture even at 20°C compared to Bowman (Fig. 6E; Supplementary Figs. S3, S4; Supplementary Tables S6, S9), comparable to the constitutively thermoresponsive long hypocotyls and petioles of Arabidopsis *elf3* mutants at 17-22°C (Jung et al., 2020; Zhu et al., 2022). These conserved architectural responses across species suggest that barley ELF3 might play a similar role in temperature responses in Arabidopsis and cereals. As cereals generally lack the thermosensory prion-like domain (Jung et al., 2020), it remains unclear whether this also refers to temperature sensing.

As especially reproductive development is vulnerable to increasing temperature and ultimately accounts for plant yields, understanding the effects of high temperature on reproductive development has become a leading research topic in barley (Fjellheim et al., 2014; Jacott and Boden, 2020). Although inflorescence development in barley is generally inhibited by high temperature (reviewed in Jacott and Boden, 2020) as observed in Bowman, transition to flowering was induced early by elevated temperatures in *elf3^BW290^* and 10_wild (Figs. 6C-E, 7A; Supplementary Fig. S4A). The observations of Bowman and *elf3^BW290^* are highly consistent with a previous report under similar conditions (Ejaz and von Korff, 2017). In contrast to the loss-of-function mutant *elf3^BW290^* with its truncated ELF3 protein due to a premature STOP codon, only a single amino acid substitution differentiates the two alleles of HIF pair 10 (W669G, Fig. 2) (Zakhrabekova et al., 2012; Zahn et al., 2022, Preprint). As transcript levels of *ELF3* remained unchanged under diurnal conditions between the two sister lines in our previous experiments (Zahn et al., 2022, Preprint), the W669G substitution may be responsible for functional differences on the protein level between the elite and wild alleles. And indeed, the W669G substitution was predicted to affect the secondary structure of ELF3 protein, putatively disturbing protein-protein interactions (Zahn et al., 2022, Preprint). Under diurnal conditions, *elf3^BW290^* promotes flowering by relieving the repression of *Ppd-H1* expression, whereas 10_wild induces the transcript levels of *FT1* and *VRN1* without influencing the transcript levels of *Ppd-H1* (Ejaz and von Korff, 2017; Zahn et al., 2022, Preprint). Surprisingly, we also observed almost no temperature effect on the transcript levels of *Ppd-H1, FT1*, and *VRN1* during inflorescence development (Fig. 7B; Supplementary Table S12). However, associated with meristem development, the transcript levels of floral inducer MADS-box genes *BM3* and *BM8* were induced by elevated temperatures exclusively in 10_wild (Fig. 7; Supplementary Table S12). On the other hand, down-regulation of *BM3* and *BM8* was reported to be correlated with late flowering in sensitive lines responding to drought (Gol et al., 2021). As under natural conditions, high temperature usually accompanies drought, these observations suggest a convergent responsive pathway. Considering the molecular connection between *ELF3* and *BM3*/*BM8*, gibberellin (GA) may be a potential mediator. It was reported that constitutive early flowering of *elf3* mutants is a result of induced GA biosynthesis, which can be blocked by the GA inhibitor paclobutrazol (Boden et al., 2014). In addition to FT1, GA synthesized in leaves might act as a mobile signal, delivering florigenic information to the shoot apex to accelerate flowering (King and Evans, 2003). Similarly, in Arabidopsis GA biosynthesis is induced by high temperatures, and it has been suggested that bioactive GA contributes to thermomorphogenesis by delivering temperature signals from the root to the shoot via one of its precursors (Stavang et al., 2009; Camut et al., 2019). These results may provide insights for future research to study the role of GA in barley temperature response.

With its multiple functions, *ELF3* has been described as one of the key plasticity genes in plants (Laitinen and Nikoloski, 2019). Natural selection of these genes leads to the phenotypic and developmental plasticity enabling acclimation responses to changing environments (Blackman, 2017). In Arabidopsis, natural variation of *ELF3* has contributed to our understanding of its role in the circadian clock and thermomorphogenesis (Anwer et al., 2014; Box et al., 2015; Raschke et al., 2015). To exploit barley natural variants of *ELF3* regarding temperature response, we selected three HIF pairs which have shown varying phenotypes between sister lines under field conditions and the exotic *ELF3* alleles of all three pairs carried W669G substitution in ELF3 (Fig. 2) (Zahn et al., 2022, Preprint). Surprisingly and in contrast to the above-described significant phenotypes of HIF pair 10, exotic *ELF3* alleles in HIF pairs 16 and 17 did not show a comparable temperature sensitivity (Figs. 3, 5; Supplementary Fig. S4; Supplementary Tables S2-9), although they carried four more amino acid substitutions in addition to the W669G replacement in ELF3 (Fig. 2). This discrepancy suggests that our observations of HIF pair 10 rely on the interaction of *ELF3* with downstream genes *Ppd-H1* and/or *VRN1*. And indeed, while HIF pair 10 carries the wild allele of *Ppd-H1*, HIF pairs 16 and 17 have the elite *Ppd-H1;* HIF pair 16 also has the wild allele of *VRN1*, whereas HIF pairs 10 and 17 contain the elite *VRN1* (Supplementary Fig. S1) (Zahn et al., 2022, Preprint). Therefore, the wild *Ppd-H1* in HIF pair 10 is likely a prerequisite for the temperature phenotypes caused by exotic *ELF3*, albeit its expression did not differ (Figs. 6C-E, 7B; Supplementary Fig. S4A). In addition, the interesting temperature-insensitive early flowering of 17_wild could result from the interaction of exotic *ELF3* and elite *Ppd-H1* (Fig. 6C, D). Furthermore, the overall late flowering of HIF pair 16 (not flowered until the end of the experiment) can be explained by the wild *VRN1* (Fig. 6C, D). These hypotheses are supported by previous reports showing allelic effects of *Ppd-H1* and *VRN1* in barley temperature response (Ejaz and von Korff, 2017; Ochagavía et al., 2021). These data emphasize the complex interactions of ELF3 with photoperiod and vernalization pathways in barley flowering control at elevated temperatures, which deserve to be studied in more detail in the future.

Besides the scientific interest in flowering time regulation, the ultimate goal of plant production under global warming conditions is to prevent yield loss at elevated temperatures. Consistent with previous reports (Dias and Lidon, 2009; Ejaz and von Korff, 2017; Ochagavía et al., 2021), we ob-served general negative effects of high temperature on several yield parameters (Fig. 8; Supplementary Fig. S4D-F). The number of grains was strongly affected by high temperature in Bowman and 10_elite (Fig. 8C). However, the reduction in grain number was not observed in *elf3^BW290^* and 10_wild, resulting in mitigated yield loss, which is potentially achieved by the nature of early flowering (Fig. 8C, F). Early flowering or early maturing is generally coupled with yield loss, due to reduced leaf area and time available for photosynthetic assimilate production, whereas it is also a strategy of plants to avoid upcoming seasonal stressful conditions such as drought and heat in summer (Shavrukov et al., 2017). Under global warming scenarios, various species flower much earlier, and this trend is predicted to continue, which indicates early flowering being an adaptive response (Parmesan and Yohe, 2003; Zheng et al., 2016; Büntgen et al., 2022). Providing further evidence, the short life cycle of *elf3^BW290^* and 10_wild in our study at elevated temperatures is also characterized by open canopy architectures and increased yield stability (Figs. 3, 5, 6C-E, 8C-F; Supplementary Fig. S3; Supplementary Tables S2-9). Our data therefore establish *ELF3* as a potentially important player contributing to barley phenotypic and developmental acclimations to elevated temperatures. While the mechanistic consequences of allelic ELF3 variants require extensive molecular studies, the presence of conditional phenotypic effects of exotic ELF3 variants in response to elevated ambient temperatures encourages systematic exploitation of this genetic resource for breeding of climate resilient crops.

## Supplementary data

**Fig. S1.** Genomic setup of the used HIFs.

**Fig. S2.** Percentage of plants with coleoptile tillers (T0).

**Fig. S3.** Effects of exotic *ELF3* alleles and elevated temperatures on barley side-view plant area and architecture.

**Fig. S4.** Principal component analysis (PCA) of traits in growth and development experiment.

**Fig. S5.** Growth and development phenotypes, and spike parameters in the yield experiment.

**Table S1.** Sequences of primers used in this study.

**Table S2.** Plant height during early growth of barley seedlings (related to Fig. 3A).

**Table S3.** Length and width of the first and second leaves (related to Fig. 3D).

**Table S4.** Plant height during barley growth and development (related to Fig. 5A).

**Table S5.** Top-view plant area during barley growth and development (related to Fig. 5B).

**Table S6.** Side-view plant area during barley growth and development (related to Fig. S2A).

**Table S7.** Plant volume during barley growth and development (related to Fig. 5C).

**Table S8.** Top-view convex hull area during barley growth and development (related to Fig. 5D).

**Table S9.** Side-view convex hull area during barley growth and development (related to Fig. S2B).

**Table S10.** Total tiller number during barley growth and development (related to Fig. 6A).

**Table S11.** Chlorophyll content in the second leaf during barley growth and development (related to Fig. 6B).

**Table S12.** Transcript levels of barley flowering genes during growth and development (related to Fig. 7B).

**Dataset S1.** Promoter sequences of *ELF3* in HIF pair 10.

**Dataset S2.** Correlation of temperature responses amongst all traits.

## Acknowledgements

We thank Tanja Zahn for generating the plant material, and Carolin Delker for providing feedback on correlation analysis. We also thank Annette Pahlich, Martin Koch, and Danny Denks for their assistance in the yield experiment and Matthias Reimers for qPCR support.

## Author contributions

KP and MQ conceived the study; ZZ and FE performed the growth and development experiment; SB performed the temperature assay on plates; ZZ and JT performed the gene expression analysis and meristem imaging; ZZ and AS performed the yield experiment; ZZ, FE, and SB sequenced *ELF3* in HIF pairs; AM performed growth curve modeling, PCA, and correlation analysis; ZZ wrote the manuscript; MQ edited the manuscript.

## Data availability statement

The data that support the findings of this study are available within the article and its supplementary data published online.

---

**Figure S1.** Genomic setup of the used HIFs. Comparison of two sister lines (upper chromosomes, elite line; lower chromosomes, wild line) in each HIF pair based on the genotype data generated from the Infinium iSelect 50k SNP chip (Zahn *et al*., 2022, Preprint). Black regions were not covered in genotyping. Green and orange parts represent homozygous elite and wild regions, respectively, whereas yellow parts represent heterozygous locus. The arrows indicate *ELF3* locus on chromosome 1H. The additional seven major flowering time genes exhibited the same fixed homozygous alleles between sister lines in all three HIF pairs. Window scaling was based on length proportion and 200 windows were created for the longest chromosome.

**Figure S2.** Percentage of plants with coleoptile tillers (T0). Bowman, *elf3^BW290^* and HIF pairs were grown in LD (16 h light/ 8 h dark) at day/night temperatures of 20/16°C (20°C). At 5 DAS, seedlings were shifted to 28/24°C (28°C) or were kept at 20°C. The formation of T0 was scored on 18 DAS (*n*=11).

**Figure S3.** Effects of exotic *ELF3* alleles and elevated temperatures on barley side-view plant area and architecture. (A, B) Bowman, *elf3^BW290^* and HIF pairs were grown in LD (16 h light/ 8 h dark) at day/night temperatures of 20/16°C (20°C). At 5 DAS, seedlings were shifted to 28/24°C (28°C) or were kept at 20°C. From 8 DAS, plants were imaged every two to four days. Side-view plant area (A) and convex hull area (B) were obtained using HTPheno and Hull & Circle pipelines (average of two side-views). Lines represent the mean and ribbons indicate SEM (*n*=11). Hashtags (for Bowman and HIF elite lines with *HvELF3*) and asterisks (for *elf3^BW290^* and HIF wild lines with *HspELF3*) indicate significant differences between two temperature treatments (# and *, *P*<0.05; ## and **, *P*<0.01; ### and ***, *P*<0.001; two-way ANOVA).

**Figure S4.** Principal component analysis (PCA) of traits in growth and development experiment. (A, B) PCA was based on the correlation matrix using the arithmetic means of all obtained traits (A), or the derived growth curve modeling traits (B) from growth and development experiment.

**Figure S5.** Growth and development phenotypes, and spike parameters in the yield experiment. Bowman, *elf3^BW290^*, and HIF pair 10 plants were grown in greenhouse conditions with LD (16 h light/ 8 h dark) and day/night temperatures of 20/16°C (20°C). Plants reached three-leaf stage were shifted to 28/24°C (28°C) or were kept at 20°C. The developmental stage (A) was scored at 73 DAS, whereas plant height (B) and total tiller number (C) were scored at 118 DAS. The developmental stage of all *elf3^BW290^* plants at both temperatures were over BBCH-59 at 73 DAS. At maturity, the spike parameters: ear length (D), number of florets per spike (E), and grain/floret ratio (F) were scored. Boxes show medians and interquartile ranges. The whiskers extend to 1.5x interquartile ranges. Dots represent values of individual plants (*n*=12-15). Different letters above or below the boxes indicate significant differences (two-way ANOVA and Tukey’s HSD test, *P*<0.05).

