## Supplementary Figures for "An exotic allele of barley *EARLY FLOWERING 3* contributes to developmental plasticity at elevated temperatures"

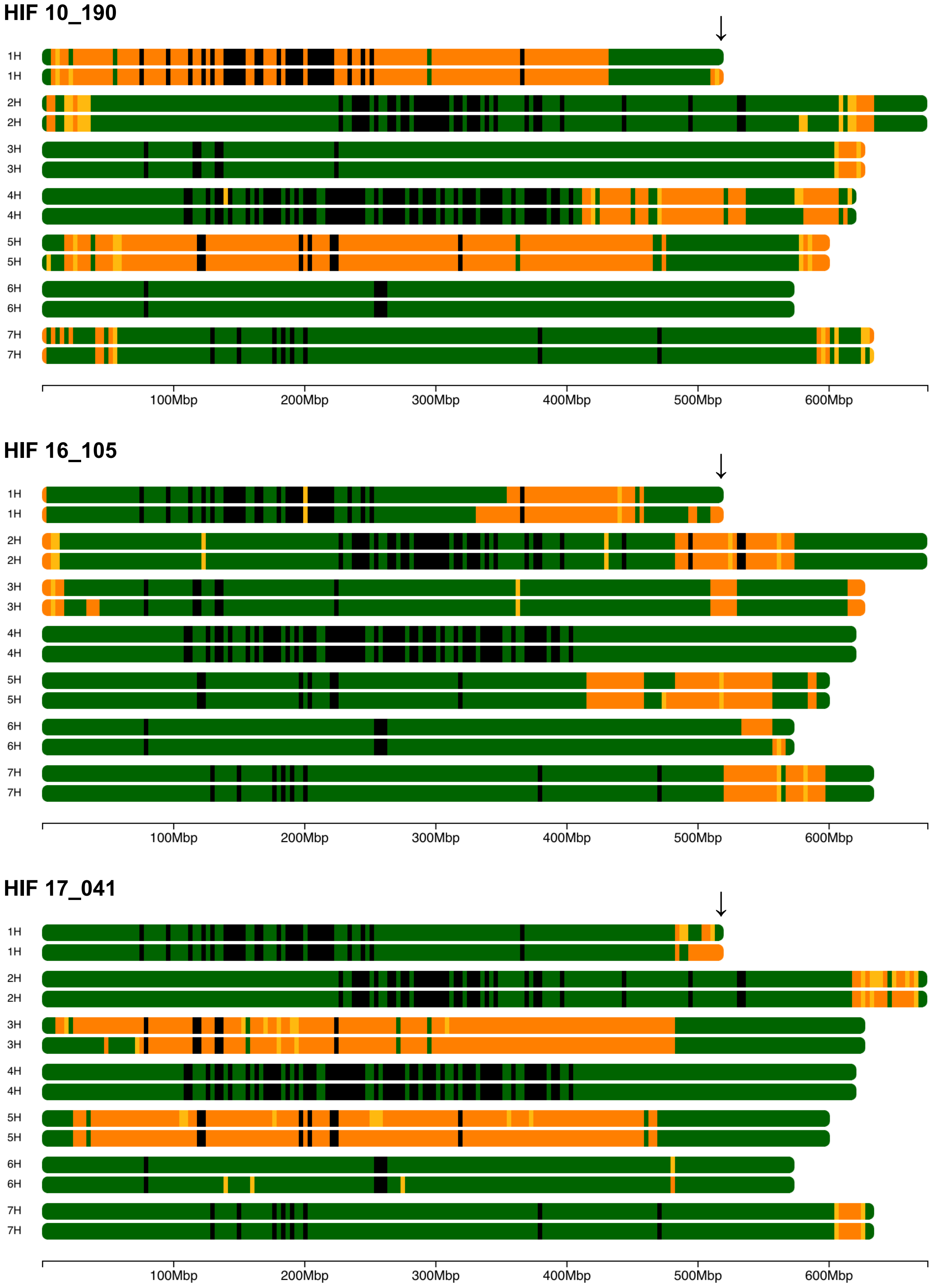


**Fig. S1.** Genomic setup of the used HIFs. Comparison of two sister lines (upper chromosomes, elite line; lower chromosomes, wild line) in each HIF pair based on the genotype data generated from the Infinium iSelect 50k SNP chip (Zahn *et al.*, 2022, Preprint). Black regions were not covered in genotyping. Green and orange parts represent homozygous elite and wild regions, respectively, whereas yellow parts represent heterozygous locus. The arrows indicate *ELF3* locus on chromosome 1H. The additional seven major flowering time genes exhibited the same fixed homozygous alleles between sister lines in all three HIF pairs. Window scaling was based on length proportion and 200 windows were created for the longest chromosome.

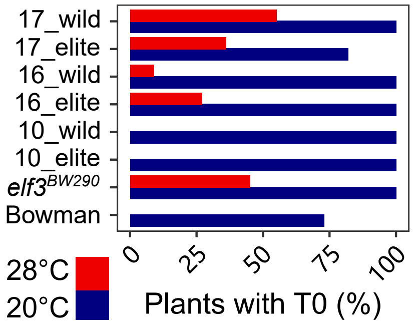


**Fig. S2.** Percentage of plants with coleoptile tillers (T0). Bowman, *elf3*$^{BW290}$ and HIF pairs were grown in LD (16 h light/ 8 h dark) at day/night temperatures of 20/16°C (20°C). At 5 DAS, seedlings were shifted to 28/24°C (28°C) or were kept at 20°C. The formation of T0 was scored on 18 DAS (*n*=11).

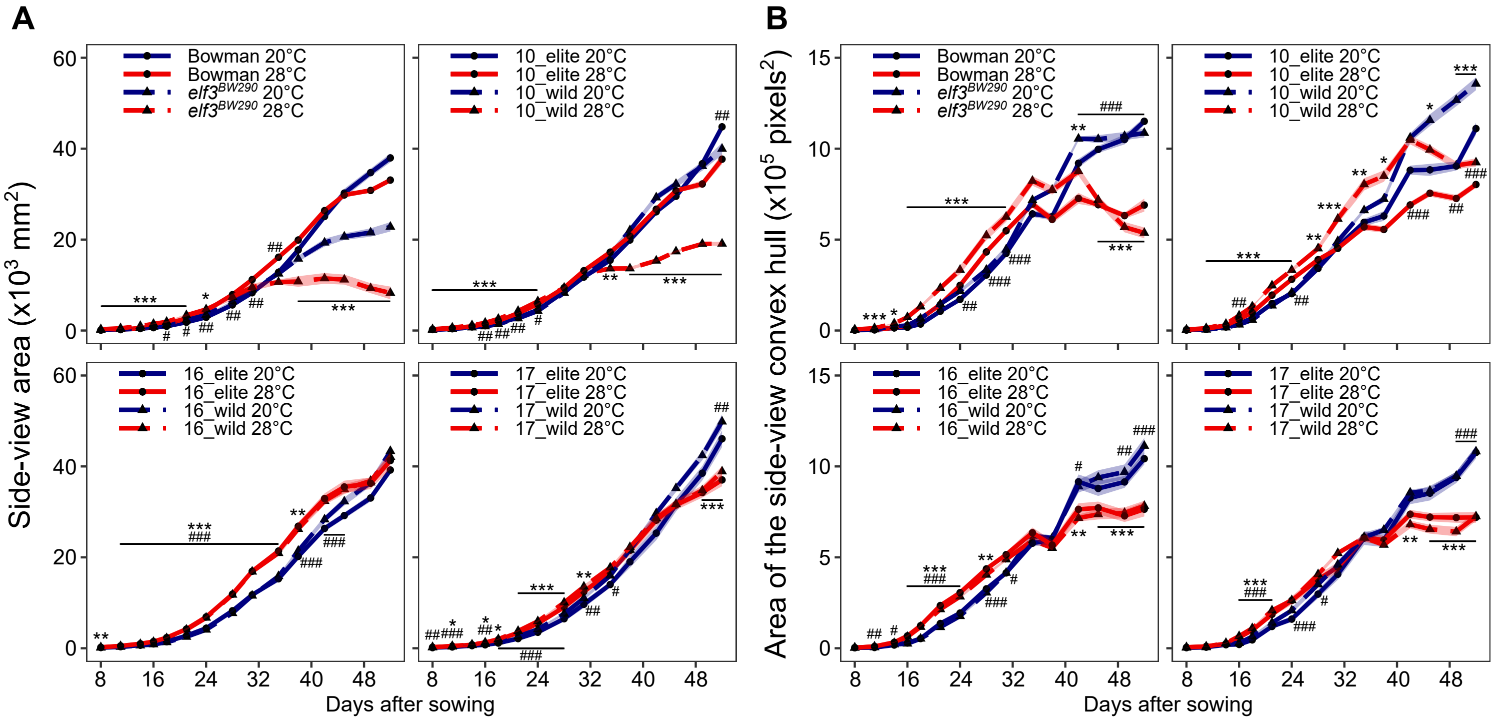


**Fig. S3.** Effects of exotic *ELF3* alleles and elevated temperatures on barley side-view plant area and architecture. (A, B) Bowman, *elf3*$^{BW290}$ and HIF pairs were grown in LD (16 h light/ 8 h dark) at day/night temperatures of 20/16°C (20°C). At 5 DAS, seedlings were shifted to 28/24°C (28°C) or were kept at 20°C. From 8 DAS, plants were imaged every two to four days. Side-view plant area (A) and convex hull area (B) were obtained using HTPheno and Hull & Circle pipelines (average of two side-views). Lines represent the mean and ribbons indicate SEM (*n*=11). Hashtags (for Bowman and HIF elite lines with *HvELF3*) and asterisks (for *elf3*$^{BW290}$ and HIF wild lines with *HspELF3*) indicate significant differences between two temperature treatments (# and *, *P*<0.05; ## and **, *P*<0.01; ### and ***, *P*<0.001; two-way ANOVA).

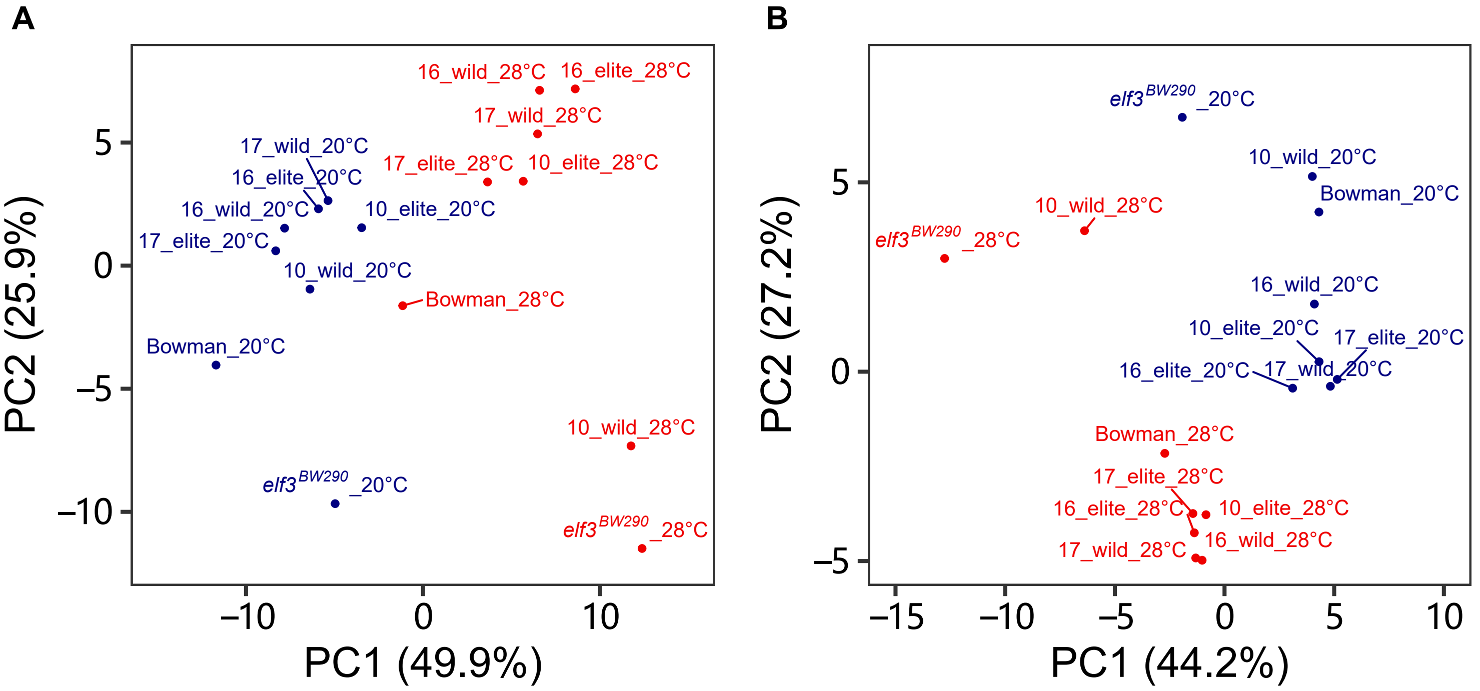


**Fig. S4.** Principal component analysis (PCA) of traits in growth and development experiment. (A, B) PCA was based on the correlation matrix using the arithmetic means of all obtained traits (A), or the derived growth curve modeling traits (B) from growth and development experiment.

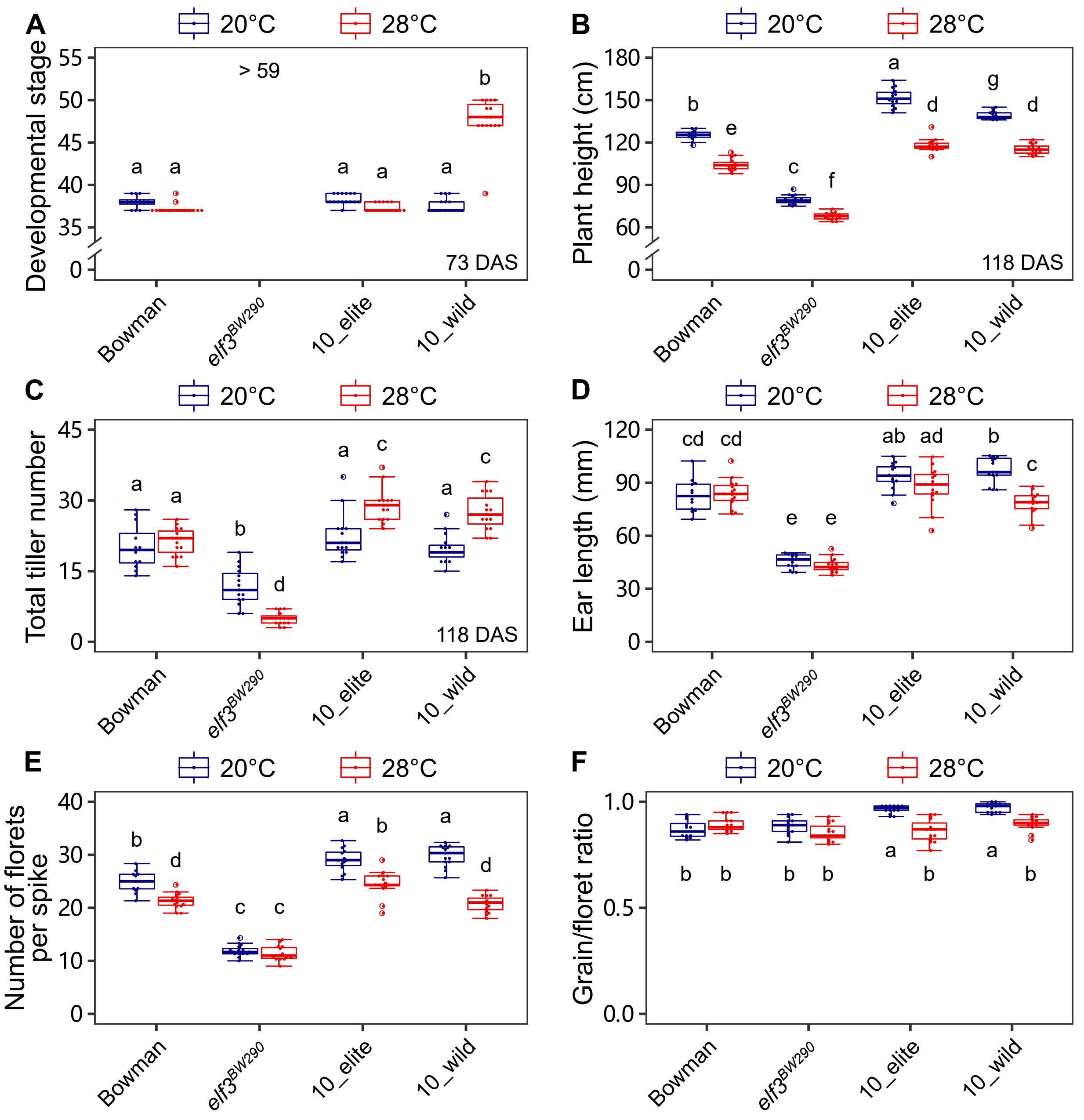


**Fig. S5.** Growth and development phenotypes, and spike parameters in the yield experiment. Bowman, *elf3*$^{BW290}$, and HIF pair 10 plants were grown in greenhouse conditions with LD (16 h light/ 8 h dark) and day/night temperatures of 20/16°C (20°C). Plants reached three-leaf stage were shifted to 28/24°C (28°C) or were kept at 20°C. The developmental stage (A) was scored at 73 DAS, whereas plant height (B) and total tiller number (C) were scored at 118 DAS. The developmental stage of all *elf3*$^{BW290}$ plants at both temperatures were over BBCH-59 at 73 DAS. At maturity, the spike parameters: ear length (D), number of florets per spike (E), and grain/floret ratio (F) were scored. Boxes show medians and interquartile ranges. The whiskers extend to 1.5x interquartile ranges. Dots represent values of individual plants ($n$=12-15). Different letters above or below the boxes indicate significant differences (two-way ANOVA and Tukey's HSD test, $P$<0.05).
